# 3D vascularized microtumors unveil aberrant ccRCC vasculature and differential sensitivity to targeted treatments

**DOI:** 10.1101/2025.03.27.645644

**Authors:** Noémie Brassard-Jollive, Yoann Atlas, Camille LM Compère, Corinne Ardidie-Robouant, Philippe Mailly, Morad El Bouchtaoui, Virginie Lelarge, Guillaume Blot, Nathalie Josseaume, Stéphanie De Oliveira, Christophe Helary, Christophe Leboeuf, Isabelle Cremer, Mathilde Sibony, Guilhem Bousquet, Stéphane Germain, Laurent Muller, Catherine Monnot

**Affiliations:** Center for Interdisciplinary Research in Biology (CIRB), College de France, CNRS UMR7241, INSERM U1050, PSL Research University, Paris, France; Sorbonne Université, Collège doctoral, F-75005 Paris, France; Université Paris Cité, INSERM, UMR_S942 MASCOT, Paris, France; Centre de recherche des Cordeliers, Universite Paris Cité, Sorbonne Université, INSERM UMRS1138, Paris, France; Laboratoire de Chimie de la Matière Condensée de Paris (LCMCP), Sorbonne Université, CNRS, F-75005 Paris, France; Hôpital Cochin, Université Paris Cité, Department of Pathology, Paris, France; Université Sorbonne Paris Nord, Villetaneuse, France; AP-HP, Hôpital Avicenne, Oncologie Médicale, Bobigny, France

**Author notes:** contributed equally to this work (Noémie Brassard-Jollive, Yoann Atlas). contributed equally to this work (Stéphane Germain, Laurent Muller, Catherine Monnot). Centre de recherche des Cordeliers, Universite Paris Cité, Sorbonne Université, INSERM UMRS1138, Paris, France (Camille LM Compère, Catherine Monnot).

**Keywords:** tumor vascularization, tumor microenvironment, 3D co-culture models, clear cell renal cell carcinoma, anti-angiogenic therapy

## Abstract

Clear cell renal cell carcinoma (ccRCC) is largely driven by Von Hippel Lindau (VHL) protein deficiency, promoting epithelial-mesenchymal transition, invasion, and hypervascularization, mediated by vascular endothelial growth factor (VEGF) signaling resulting in a structurally abnormal capillary network, which remains insufficiently defined. Previous studies correlating patient outcome with microvascular density yielded diverse results, underscoring the limitations of conventional parameters to fully capture vascular complexity. While VEGF-targeted anti-angiogenic first-line therapies such as sunitinib prolong progression-free survival in metastatic ccRCC, their efficacy is hampered by resistance mechanisms.

This study aims to elucidate the three-dimensional architecture of ccRCC-specific vasculature in the tumor microenvironment, and its response to targeted therapies. Analysis of human ccRCC samples identified two distinct vascular structures, markedly differing from tumor capillaries, termed ponds and sheets, which were further characterized in patient-derived xenografts using advanced 3D microscopy on optically cleared samples. Ponds are large, dilated, irregular structures with wide cavity, whereas sheets are thin, elongated, and collapsed structures. To further dissect endothelial network morphogenesis, we developed an innovative *in vitro* 3D vascularized microtumor model, faithfully recapitulating the aberrant pond architecture. Dynamic live imaging unraveled the temporal relationship between tumor invasion and pond morphogenesis. Additionally, drug sensitivity assays demonstrated that ponds exhibit lower responsiveness to sunitinib compared to tumor capillaries.

Altogether, our 3D model not only captures a specific architecture of ccRCC vascular network but also provides mechanistic insight into its development and therapeutic sensitivity. This model offers a promising avenue for personalized treatment assessment and for identification of novel therapeutic strategies.

**Statement of significance:** A co-culture-engineered model integrating tumor spheroid invasion and capillary morphogenesis recapitulates the ccRCC endothelial structures and their response to treatments.

## INTRODUCTION

Clear cell renal cell carcinoma (ccRCC) is the most prevalent form of renal carcinoma, accounting for approximately 70% of cases^1^. This aggressive cancer, often spreading hematogenously, exhibits a high metastatic potential and poor prognosis, largely driven by von Hippel-Lindau (*VHL*) gene alterations in about 80% of cases. These mutations stabilize hypoxia-inducible factors (HIFs) in tumor cells, triggering transcriptional programs that promote tumor invasion, epithelial-to-mesenchymal transition (EMT), angiogenesis, and vascular remodeling. This includes the upregulation of key genes coding for vascular endothelial growth factor (VEGF) and hepatocyte growth factor (HGF) receptor tyrosine kinase c-MET^1,2^. Consequently, this HIF-driven pseudo-hypoxia induces hypervascularization^3^, characterized by a chaotic vascular network exhibiting loops, dead-ends, tortuous capillaries highly variable in size, shape and connectivity^4^. The tumor microenvironment (TME) of ccRCC is thus characterized by high levels of CD31^+^ or CD34^+^ endothelial signals, revealed by elevated intratumoral microvessel density (MVD), and by structurally abnormal capillary network. However, MVD’s prognostic value is matter of debate, as studies report inverse^5,6^ or positive^7,8^ correlations with survival. Growing evidence suggests that discrepancies in MVD and prognostic value may arise out of the architectural complexity of the tumor vasculature rather than vessel density alone. In support of this, microvessel fractal dimension, a parameter quantifying vascular complexity, has been identified as a prognostic marker positively correlated with patient survival in renal cell carcinoma^9^. Furthermore, morphometric analyses have identified two distinct vascular phenotypes in ccRCC, providing additional prognostic information beyond MVD alone^8^.

Characterizing the 3D vascular architecture within the ccRCC microenvironment presents a major challenge. The TME consists of heterogeneous and interacting cancer and stromal cells, such as endothelial cells, fibroblasts and immune cells, embedded within a remodeled extracellular matrix (ECM)^10^. The pseudo-hypoxic niche facilitates tumor cell invasion and plays a major role in malignant progression^11^. Collagen I, a key driver of cancer cell invasion^12^, is predominantly produced by fibroblasts in ccRCC^13–15^.

Targeting the VEGF pathway has been the cornerstone of metastatic ccRCC therapy for decades^1,16^, with multi-tyrosine kinase inhibitors (TKI) like sunitinib improving survival and standing as reference in clinical trials^17^. However, resistance to sunitinib often resulted in post-treatment tumor “flare-ups” of tumor invasiveness and dissemination^18^. Alternative therapies targeting mTOR or c-MET have shown limited efficacy compared to TKI^2,19,20^. Alternative treatments, targeting mammalian target of rapamycin (mTOR) or HGF pathways have been evaluated in clinical trials^16,19^. Temsirolimus is an inhibitor of mTOR playing a critical role in regulating cellular protein synthesis and the translation of HIF-1α. However, multiple clinical trials comparing mTOR inhibitors to sunitinib have shown superior efficacy for the TKI^20^. Crizotinib, an inhibitor of c-MET demonstrated moderate anti-angiogenic and anti-tumor activity compared to sunitinib in both TKI-sensitive and resistant models^2^.

With recent treatment strategies combining TKIs and immunotherapy in ccRCC^16^, deciphering the impact of anti-angiogenic therapies on vascularization and invasion is crucial. A better understanding of the 3D architecture of the vascular network requires to analyze the functional and dynamic interaction between angiogenesis and TME, key mediator of the therapeutic resistance^19^.

This study aimed to develop an innovative and relevant 3D *in vitro* model that mimics the ccRCC vasculature, to evaluate its response to current and emerging therapeutic strategies^21^. We recreated the TME by embedding both tumor and endothelial cells within a collagen I hydrogel and culturing them in fibroblast-conditioned medium. Using the human invasive VHL-deficient cell line RCC4, known for its tumorigenic and angiogenic capacities^22^, or its epithelial-like, VHL-restored counterpart, RCC4-VHL^23^, along with primary cultures of human umbilical vein endothelial cells (HUVECs), the model allowed the analysis of tumor invasion and capillary morphogenesis. Notably, through 3D morphological analysis, we identified and characterized endothelial structures, termed *ponds*, observed in patient-ccRCC samples and patient-derived xenografts (PDX). These structures exhibited reduced sensitivity to sunitinib compared to tumor capillaries, highlighting their potential role in therapeutic resistance. This model provides novel insights into ccRCC vascular heterogeneity and may contribute to the development of more effective anti-angiogenic strategies.

## MATERIALS AND METHODS

### Staining of human clear cell renal cell carcinoma and patient-derived xenograft samples

Slices of six human samples of clear cell renal cell carcinomas were analyzed after fixation in 10% formalin, paraffin embedding and immunostaining for CD31, performed in the department of pathology of Cochin’s hospital. Sample collection for further research analysis was approved by the local ethics committee of CARPEM (CAncer Research for Personalized Medicine) from the integrated cancer research site (SIRIC) accredited by the French National Cancer Institute (INCa). Subcutaneous engraftment of human primary ccRCC fragment samples in nu/nu athymic mice, tumor growth and CD31 immunostaining were performed for four PDX samples, as previously described^24,25^. For 3D reconstruction, PDX were fixed with 4% paraformaldehyde solution and sectioned into 500 µm-slices using vibratome (HM 650 V, Leica Biosystems, Wetzlar, Germany). Slices were washed 3 times with Dulbecco’s Phosphate Buffered Saline 1X (PBS, ThermoFisher Scientific, Waltham, MA, USA) and permeabilized for 2 days in PBS containing 2% triton-X-100 (Sigma-Aldrich, Saint-Louis, Mo, USA). Immunofluorescent stainings were performed in PBS-Triton-X-100 0.01%. Vascular structures were stained with rat anti-mouse CD31 monoclonal antibody (mAb, BD Biosciences, Franklin Lakes, NJ, USA, #550274, 1/100) for 1 day. Nuclei were stained with 4′,6-diamidino-2-phenylindole (DAPI) (Invitrogen, Carlsbad, CA, USA). Stained 500 µm-slices were then clarified using RapiClear®1.52 solution (SunJin Lab, Taïwan) before imaging. Immunofluorescent staining of the vascular basement membrane was performed on 100 µm-slices of PDX samples using rabbit anti-mouse Collagen IV Ab (Novotec, Bron, France, #20451, 1/100).

### Cell lines and primary cultures

RCC4 cell line derived from a primary ccRCC tumor harboring sporadic *VHL* mutation (RCC4) and its counterpart, with restored wild-type VHL (RCC4-VHL), were a gift from Patrick H. Maxwell (Cambridge Institute for Medical, UK)^23^. RCC4-GFP and RCC4-VHL-LAR cells have been transduced and cloned respectively for stable expression of cytosolic green fluorescent protein (GFP) inserted with pLenti vector (ThermoFisher Scientific, Whaltham MA, USA) and fluorescent reporter of filamentous actin (LifeAct Ruby, LAR). Both cell lines RCC4 and RCC4-VHL have been transduced overnight with lentiviral vectors. Homogeneous fluorescent cell populations have been screened by fluorescent activated cell sorting analysis based on middle fluorescence intensity. Two clones from single-cell cultures have been selected through analysis of 2D- and 3D-cell behavior (RCC4-GFP-1, RCC4-GFP-2; RCC4-VHL-LAR-1, RCC4-VHL-LAR-2). Cells were cultured in Dulbecco’s Modified Eagle Medium (DMEM, Invitrogen) supplemented with 10% fetal bovine serum (FBS, Invitrogen). Primary culture of adult normal human dermal fibroblasts (NHDF, Promocell, Heidelberg, Germany) were grown in fibroblast growth medium 2 (FGM2, PromoCell). Conditioned media from RCC4 (RCC4-CM), RCC4-VHL (RCC4-VHL-CM) and fibroblast (NHDF-CM) cells were obtained from confluent cultures in ECGM2 depleted for VEGF_165_ (hereafter called VEGF) (ECGM2-ΔVEGF) and collected every day between 2 to 10 days. Primary culture of human umbilical vein endothelial cells (HUVECs) were isolated from cords provided by AP-HP, Hôpital Saint-Louis, Unité de Thérapie Cellulaire, CRB-Banque de Sang de Cordon, Paris, France (authorization number AC-2016-2759) and cultured as previously described^26^ in endothelial cell growth medium 2 (ECGM2, PromoCell). Experiments were performed between passages 1 and 5 for primary cell cultures.

### Immunoblotting and ELISA assay

Confluent ccRCC cell lines grown for 3 days were lysed in a buffer containing 25 mM Tris pH 7.5, 100 mM NaCl, 5 mM EDTA, 0.5% DOC, 0.5% NP40, and a protease inhibitor cocktail. The lysates were centrifuged for 30 minutes at 15,000g. Supernatants were treated with Laemmli buffer. Proteins were separated by SDS-PAGE and transferred to PVDF membranes. The following primary antibodies were used: mouse anti-human SNAIL mAb (Merck Millipore, Burlington, MA, USA, #MABE167, 1/1000), rabbit anti-human LOX mAb (Abcam, Cambridge, UK, #Ab174316, 1/1000), rabbit anti-human LOXL2 mAb (Cell Signaling Technology, Danvers, MA, USA, #69301, 1/500) and rabbit anti-human actin polyclonal Ab (Abcam, #Ab8227, 1/5000). Immunodetection was performed with alkaline phosphatase– conjugated secondary antibodies (Jackson ImmunoResearch Europe, Cambridgeshire, UK), and Pierce ECL Western substrate (ThermoFisher Scientific, Waltham, MA, USA). For conditioned media collected under similar culture conditions, samples were centrifuged at 4°C for 10 minutes at 900 rpm to eliminate cellular debris. The VEGF concentration was determined by ELISA according to the manufacturer’s instructions (R&D Systems, Minneapolis, MN, USA) and normalized to the total protein content of the cells. The experiment was performed in duplicate.

### Collective migration and proliferation assays

Collective migration of ccRCC cell lines or HUVECs was monitored using a live-cell imaging system (IncuCyte Zoom, Essen Bioscience, United States). For the assay, 15,000 RCC4 cells or 20,000 HUVECs were seeded per well in a 96-well plate (Corning, NY, USA) containing Oris stoppers (Platypus Technologies, Madison, WI, USA). Stoppers were removed the day after and complete growth medium was replaced by 2% FBS-containing medium for ccRCC cell lines or by ccRCC cell line-CM for HUVECs. The migration assay was monitored for 24 hours, with images captured every 6 hours for ccRCC cell lines and every 2 hours for HUVECs, at 4x magnification. Data analysis was performed using IncuCyte analysis software, to quantify cell surface area coverage as confluence values. Migration rates were calculated for ccRCC cell lines or HUVECs, using IncuCyte analysis software and ImageJ respectively. The proliferation of ccRCC cell lines was monitored using IncuCyte Zoom. Cells (1,000 cells/well) were seeded in a 96 well-plate (Corning) in DMEM supplemented with 7.5% FBS. The assay was monitored for 72 hours, with image acquisition every 3 hours at 20x magnification. Data analysis was performed using IncuCyte analysis software, for quantification of the cell surface area coverage as confluence values. The growth curve f(t)=ln(confluence mask %) was drawn, and a linear regression was established. The generation time (G), defined as the time required for cells to divide, was calculated using the slope (µexpo) of the linear regression associated with G = ln(2)/µexpo formula.

### Spheroid formation and invasion assay

Spheroids were formed using the hanging drop assay. Cells were seeded at densities of 250, 500 or 1,000 cells per 25 µL drop of DMEM medium containing 0.06% methylcellulose (Sigma-Aldrich) and supplemented with 17.5% FBS (Invitrogen). The drops were placed on the inner side of a petri dish lid, which was then inverted and positioned over a petri dish filled with PBS containing 1% FBS, and placed into incubator during 48 hours.

Spheroid invasion assays were performed using the fluorescent clones RCC4-GFP and RCC4-VHL-LAR. Spheroids (500 cells each) were embedded in a 2 mg/mL collagen I solution, prepared in-house as previously described^27^, mixed with 10x M199 solution (Sigma-Aldrich) buffered at pH 7 at 20°C using a NaHCO3 solution. The embedded spheroids were cultured in 96-well phenoplates (Revvity, Waltham, MA, USA) for 3 days in various media: DMEM; ECGM2 depleted of epidermal growth factor (EGF), basic fibroblast growth factor (bFGF), insulin-like growth factor (IGF), VEGF and hydrocortisone (ECGM2-Δ5); ECGM2 containing EGF (ECGM2-Δ5+EGF); ECGM2 depleted only of VEGF (ECGM2-ΔVEGF); or in NHDF-CM.

### Capillary formation and stabilization assays

3D angiogenesis assays were performed using two different approaches, with conditioned media being replaced daily in both models. In the first approach, HUVECs were encapsulated at a density of 1.5×10^6^ cells/mL in a 2 mg/mL collagen I hydrogel (pH 7), as previously described^28^. Cells were cultured for 5 days in presence of various cell-derived conditioned media. For the capillary destabilization assay, pre-formed capillary networks, generated over 5 days in the presence of NHDF-CM supplemented with VEGF (10 ng/mL), were further cultured for 4 days in the presence of different cell-derived conditioned media. The second approach consisted in angiogenesis assay from Cytodex beads, as previously described^26^. Capillary sprouts were cultured for 4 days in the presence of different cell-derived CM.

### Vascularized spheroid model

For the vascularized microtumor model, HUVECs were co-encapsulated at a density of 1.5×10^6^ cells/mL with RCC4 or RCC4-VHL spheroids (500 cells each) in 2 mg/mL collagen I hydrogel (pH 6.5). Cells were cultured for 4 to 7 days in the presence of NHDF-CM. For the co-culture model of cells scattered in the hydrogel, HUVECs were co-encapsulated at a density of 1.5×10^6^ cells/mL in presence of various densities of RCC4 cells (stated in the figure legend) in 2 mg/mL collagen I hydrogel (pH 6.5).

For kinetic analysis, fluorescent HUVEC-LAR or HUVEC-GFP were co-encapsulated in collagen I (2 mg/mL) hydrogel at a density of 1.5×10^6^ cells/mL, along with RCC4-GFP or RCC4-VHL-LAR spheroids (500 cells each), respectively. Cells were cultured for 6 days in presence of NHDF-CM, in similar conditions than described above. Time-lapse confocal imaging of the same spheroid was performed every 10-12 hours.

To assess the effects of drugs on tumor invasion, capillary or pond formation, RCC4 spheroids and HUVECs, either alone or in co-culture were treated with vehicle (dimethyl sulfoxide, DMSO), temsirolimus (ThermoFisher Scientific, #J63654.MB; 5, 10 or 25 µM, crizotinib (Merck Millipore, #PZ0191; 0.05 or 0.5 µM) or sunitinib (Merck Millipore, #1642358; 0.05, 0.5 or 5 µM). Drugs were added to the cell-containing hydrogels before polymerization, and the medium containing the same drugs was refreshed after one day of culture. Hydrogels were fixed following 2 days of treatment. To assess the sensitivity of pre-existing structures, vascularization of spheroids was promoted for 5 days in the presence of NHDF-CM after which treatment with sunitinib (0.05 or 0.5 µM) was applied for 2 days.

### Staining of 2D and 3D *in vitro* models

Tumor cells were seeded at a low density in an 8-wells plate coverslips (Ibidi GmbH, Gräfelfing Germany) uncoated or coated with a 37.7µg/mL solution of rat tail collagen I (Corning). After 2 days of culture, cells were fixed with 4% PFA for 10 minutes, washed 3 times with PBS and permeabilized for 20 minutes in PBS containing 0.5% triton-X-100. Immunostaining and fluorescent labeling were performed as described above.

Spheroids and cells embedded in hydrogels were fixed with 4% paraformaldehyde for 30 minutes, washed 3 times with PBS and permeabilized for 30 minutes in PBS containing 0.5% triton-X-100 (Sigma-Aldrich). Fluorescent stainings were performed in PBS containing 0.01% Triton-X-100. Actin was stained with phalloidin conjugated to Alexa Fluor −488 or −555 (Invitrogen, #A12379 or #A34055, 1/2000). Nuclei were stained with DAPI (Invitrogen). In the vascularized microtumor model, HUVECs were immunostained with mouse anti-human CD31 mAb conjugated to Alexa Fluor 647 (Abcam, #Ab215912, 1/200) and basement membrane with rabbit anti-human collagen IV Ab (Novotec, #20411, 1/80) followed by secondary anti-rabbit Ab coupled to Alexa Fluor 555 (Invitrogen, #A-31572, 1/500).

### Microscopic imaging and quantitative image analysis

High-resolution images of 500 µm-slices of PDX samples were acquired using a Zeiss (Oberkochen, Germany) LSM 980 microscope with Airyscan 2 (63x/1.1 objective). Tiled images were stitched using Zen Microscopy software (Zeiss), and 3D reconstructions were performed with Imaris software.

The formation of RCC4 or RCC4-VHL spheroids was monitored with a live-cell imaging system (EVOS M7000, ThermoFisher Scientific). Fluorescent RCC4 or RCC4-VHL cells (500 cells per well) were seeded in Nuncleon sphere-treated, U-shaped 96-well plates (ThermoFisher Scientific) in complete DMEM. Fluorescent and transmitted-light images were acquired every 15 minutes during the first 3 hours and every 45 minutes during the subsequent 6 hours. Brightfield images of RCC4 or RCC4-VHL spheroids in hanging drops were acquired by a Leica DMi1 microscope with an MC170HD camera (10x/0.3 objective).

Fluorescent images of RCC4 or RCC4-VHL spheroids and vascularized microtumors in hydrogels were captured with a Zeiss Observer Apotome 2 (10x/0.45 or 40x/1.1 objective) or a Zeiss Spinning-Disk CSU-W1 (25x/0.8 or 40x/1.1 objective). Capillary network formation and stabilization in hydrogels were studied using the Zeiss Observer Apotome 2 (10x/0.45 or 40x/1.1 objective).

Spheroid (area, perimeter) and vascular (area) parameters were measured by 2D analysis of maximal z-stack projection masks using Fiji (ImageJ software). Masks were obtained using a default threshold and particles were analyzed for a minimum pixel size of 100µm². The spheroid circularity index was calculated as 4 × π × (area/perimeter×2). Capillary parameters (length, number, branch points) were measured by 3D quantification of the network as previously described^28^. Connectivity index was adapted from the graph theory and calculated as (branch number – skeleton number)/ branch number.

### Statistical analysis

Statistical analyses were performed using Prism software version 8.0 (GraphPad Prism). Data were obtained from at least three independent experiments. Numerical variables were expressed as the mean ± SEM (standard error of the mean). The normality of the distribution was assessed using the D’Agostino-Pearson and Shapiro-Wilk tests. Based on normally distributed data and homogeneity of variance, one-way or Brown-Forsythe or Welch ANOVA tests was conducted for comparing three or more groups. If the distribution of at least one group did not meet normality or the sample size was insufficient for testing, comparisons between groups were made using the nonparametric Kruskal-Wallis test. Mann-Whitney test was conducted for comparing two groups. Differences were considered significant at p<0.05.

## RESULTS

### The ccRCC vascular network exhibits aberrant endothelial structures

Beyond the well-known exuberant vascularization, we analyzed human ccRCC samples to characterize the architectural complexity of the blood vasculature using CD31 staining. The intricate and dense CD31^+^ network (**Figure 1Aa-B**) was embedded within tumor cell masses, identifiable by their clear and extensive cytoplasm. This network included numerous tumor capillaries exhibiting a circular shape and lumens ranging from 10 to 20 µm in diameter (**Figure 1Ab**), alongside aberrant structures that markedly differed in shapes and dimensions, exhibiting two distinct architectures. The first type, termed *pond*, consisted of dilated structure displaying wide, irregular shape and large cavity exceeding 100 µm in diameter (**Figure 1Ac**). The second type, termed *sheet*, consisted of thin, elongated, collapsed structure extending continuously for over 100 µm, occasionally interconnecting with ponds (**Figure 1B**). The abundance and distribution of these aberrant vascular structures varied both intra-tumorally and inter-patient (**Figure 1A-B**).

**Figure 1.**
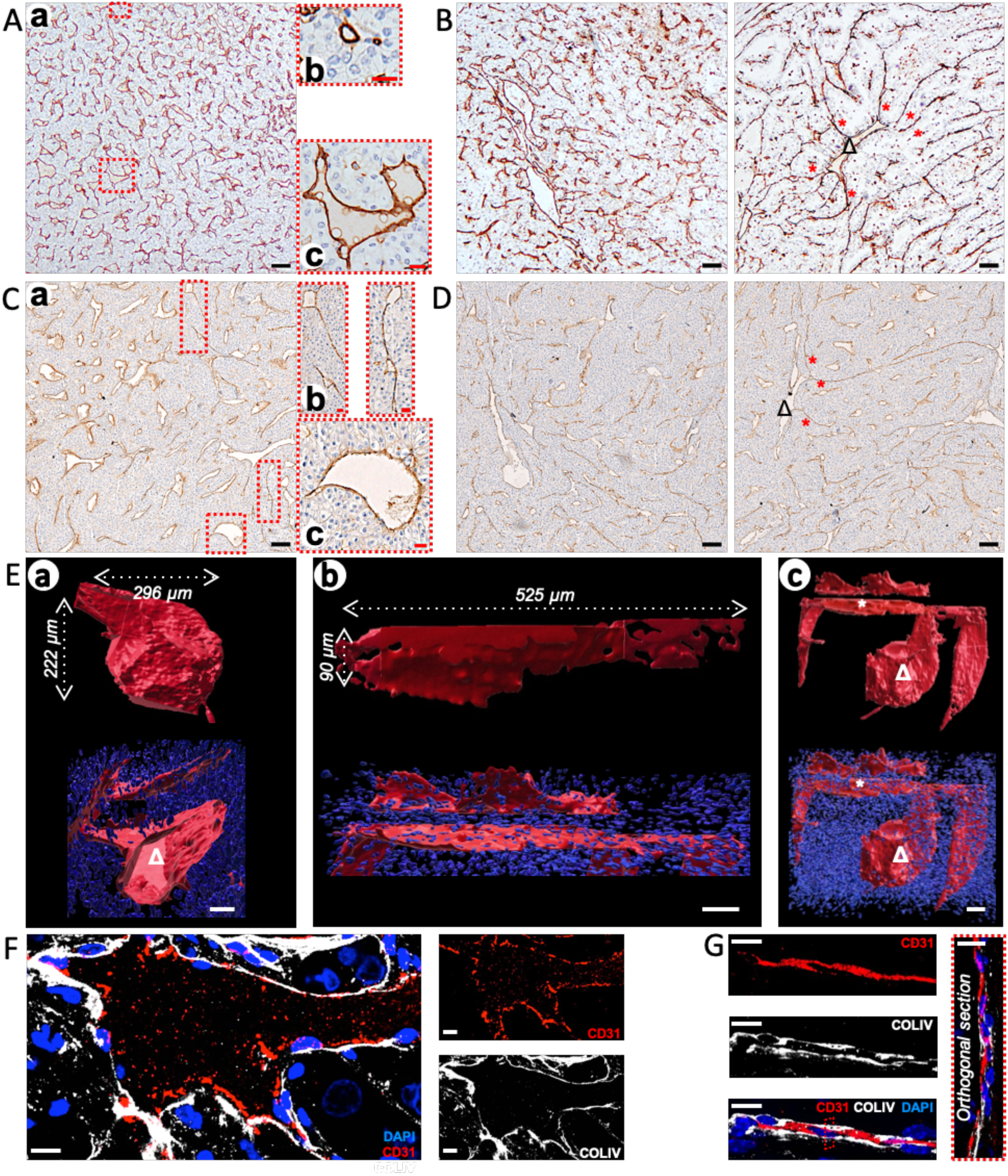
ccRCC blood vasculature contains aberrant endothelial structures in human primary tumor and in patient-derived xenograft in mouse. **A** and **B**: Immunohistochemistry for CD31 staining of two representative ccRCC tumors. Magnifications of red-dotted frames displaying blood capillary (**A, b**) and pond (**A, c**). Scale bar: 100 µm (black) or 20 µm (red). **C** and **D**: Immunohistochemistry for CD31 staining in a representative ccRCC-derived xenograft implanted in nude mouse. Magnifications of red-dotted frames displaying interconnected aberrant structures (**B, b**) and pond (**B, c**). Interconnected sheets and pond are identified by red asterisk and black capital delta. Scale bar: 100 µm (black) or 20 µm (red). **E**: 500 µm-thick section of the ccRCC-derived xenograft immunostained for CD31 (red), stained by DAPI for nuclei (blue) and clarified for imaging. Treatment with Imaris software for surface rendering and 3D reconstruction were performed on representative images of pond (**E, a**), sheets (**E, b**) and large view covering the two aberrant structures (**E, c**). Dimensions of each aberrant structure are mentioned in white (**E, a** and **b**, top panels). Clipping plane of the 3D reconstruction reveals pond lumen (**E, a**, bottom panel). Interconnected sheet and pond are identified by white asterisk and capital delta. Scale bar: 50 µm. **F** and **G**: 100 µm-thick section of ccRCC-derived xenograft immunostained for collagen IV (white), CD31 (red) and stained by DAPI for nuclei (blue) revealing pond (**F**) and sheet (**G**). Orthogonal section (y-z) was imaged in the red-dotted frame (**G**, right panel). Scale bar: 10 µm.

PDX model provided additional insights into the architectural features of the aberrant vascular structures (**Figure 1C-D**). Our findings revealed that this model recapitulated the complex vascular network observed in human samples, in line with previous reports describing the preservation of the molecular, genetic, and histopathological features of ccRCC in PDX models^29,30^. The dense tumor mass within the PDX contained numerous CD31^+^ structures with diverse morphologies, including long, thin (**Figure 1Ca-b**) or dilated, irregular (**Figure 1Ca-c**) structures, closely resembling the sheets and ponds observed in human ccRCC samples, in terms of dimension, shape, and interconnections. We hypothesized that the high volumes of ponds and sheets in the tumor mass might explain the detection of continuous CD31^+^ structures spanning hundreds of micrometers in both human and PDX samples, as seen in thin-section imaging. We thus conducted innovative 3D morphometric analyses on 500 µm-thick sections of PDX, stained for CD31 and nuclei, followed by tissue clearing. High-resolution image acquisition and stitching enabled 3D reconstruction of the irregularly dilated structures containing a cavity, extending over 200 µm in both width and height (**Figure 1Ea**) while sheets appeared as elongated, thin, and collapsed structures extending over 500 µm (**Figure 1Eb**). A large-scale view from the stitched images depicted a complex vascular network, embedded within regions of high nuclear density, consisting of two parallel sheets, one of which was connected to two distinct ponds (**Figure 1Ec** and **Supplementary Movie S1**). Additionally, a nearby endothelial capillary strongly differed from this aberrant network (**Supplementary Movie S1**). A z-stack imaging of interconnected pond and sheets within the tumor mass revealed a high density of red blood cells, densely packed within the pond cavity and forming aggregates at the pond-sheet junction (**Supplementary Movie S2** and **Figure S1**). Immunofluorescent co-staining for CD31 and the basement membrane component, collagen IV in PDX sections (**Figure 1F-G**) revealed an endothelial monolayer delineating the large pond cavity, bordered by the basement membrane on its basal side (**Figure 1F**), revealing a polarized structure. In contrast, sheets showed collagen IV on both sides of a continuous endothelium, suggesting collapsed capillaries (**Figure 1G**). In conclusion, we identified aberrant architecture of vascular structures, within the ccRCC tumoral area, differing from those of tumor capillaries and characterized by outlier dimensions and unique 3D structural patterns.

### RCC4 spheroids retain tumor-like features in a 3D model

The preservation of the ccRCC vascular network architecture in the PDX model suggests a significant impact of human tumor cells on the mouse vascular compartment, highlighting the critical role of tumor-endothelial cells interactions in shaping aberrant vasculature. Given the need to elucidate its formation and response to TKI treatments, the limitations of the PDX model including technical challenges, labor-intensity, time constraints, and high costs, despite its invaluable contribution, became evident. To overcome these limitations and enable cellular-scale analyses, we developed a 3D *in vitro* co-culture model incorporating tumor spheroids and capillaries. We first established and characterized spheroids of human ccRCC cell line RCC4 and its counterpart, with restored wild-type *VHL* gene (RCC4-VHL). Fluorescent clones of RCC4 cells expressing GFP (RCC4-GFP) and RCC4-VHL cells expressing LAR (RCC4-VHL-LAR) were generated and validated, maintaining the parental cell line features, including EMT markers, cell behavior, proliferation and migration in 2D cultures (**Supplementary Figure S2**). Immunoblot analysis confirmed the high expression of lysyl oxidase (LOX) and lysyl oxidase-like 2 (LOXL2), two hypoxia-targeted ECM-modifying enzymes expressed in ccRCC^31,32^, in RCC4 or RCC4-VHL under normoxic and hypoxic conditions or hypoxic condition only, respectively. Furthermore, in normoxia, LOXL2 followed the secretory pathway, accumulating in perinuclear organelles of RCC4 cells (**Supplementary Figure S2A-E**). In 2D cultures, RCC4 cells and their clones displayed an isolated and homogeneous pattern, whereas RCC4-VHL cells and their clones formed cohesive, epithelium-like (**Supplementary Figure S2F-G**). While generation times were comparable between RCC4 and RCC4-VHL cell lines and clones, RCC4 cells and their clones exhibited significantly higher collective migration than RCC4-VHL cells and their clones (**Supplementary Figure S2H-J**). Given their relevance as investigative tools, fluorescent clones were further analyzed in 3D conditions.

The early stages of RCC4 or RCC4-VHL spheroid formation were monitored using transmitted-light and fluorescent live-cell imaging immediately after seeding in U-shaped wells, for 9 hours (**Supplementary Movies S3-S6**). Aggregation of single cells into spheroid exhibited comparable clustering dynamics. After 48 hours, fully formed spheroids were embedded in a collagen I hydrogel (**Figure 2A**). While both cell lines initially formed morphologically similar spheroids (0 hour), their structure rapidly diverged (after 6 hours), RCC4 spheroids exhibiting numerous actin-rich protrusions, indicative of active cellular dynamics whereas RCC4-VHL spheroids underwent a compaction process.

**Figure 2.**
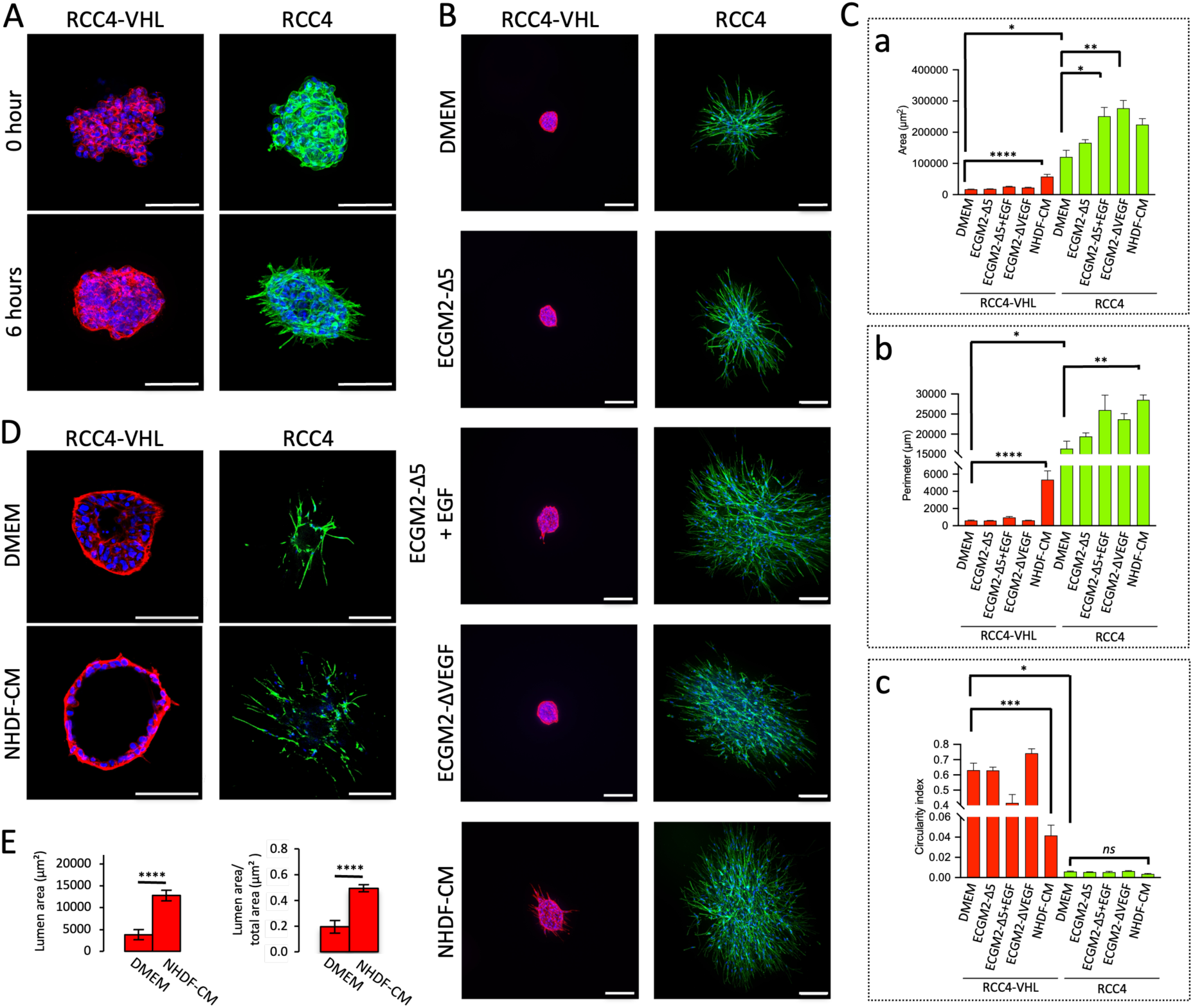
RCC4 spheroids display tumor-like features linked to the *VHL* status in 3D models. **A, B** and **D**: Spheroids of 500 RCC4-VHL or RCC4 cells embedded in collagen I hydrogel and cultured for 0 or 6 hours in complete DMEM (**A**) or for 3 days in complete DMEM or fibroblast-conditioned medium (NHDF-CM) (**B** and **D**), growth factor-depleted ECGM2 (ECGM2-Δ5) or supplemented with EGF (ECGM2-Δ5+EGF), VEGF-depleted ECGM2 (ECGM2-ΔVEGF) (**B**), stained for F-actin by phalloidin (red for RCC4-VHL, green for RCC4) and for nuclei by DAPI (blue). Images are displayed as z-stack projection (**A** and **B**) and optical section localized at the spheroid center determined by the highest diameter size (**D**). Scale bar: 100 µm (**A**) or 200 µm (**B** and **D**). **C**: Area (**a**), perimeter (**b**) and circularity index (**c**) of spheroids cultured in conditions described in panels B and D. Quantifications were performed on z-stack projections. **E**: Lumen area (left panel) and lumen area over total spheroid area (right panel) of RCC4-VHL spheroids cultured in conditions described in panels B and D. Quantifications were performed on central optical section. Graphs represent the mean of 3 independent experiments +/- SEM (**C** and **E**). Statistical analyses were performed using Kruskal-Wallis (**C**) or Welch’s t (**E**) tests. ns p>0.05, * p<0.05; ** p<0.01; *** p<0.001; **** p<0.0001.

To optimize the 3D co-culture model that mimicked tumor characteristics, it was essential to identify a medium compatible with capillary growth. Endothelial cells alone are unable to form a stable *in vitro* microvascular network of polarized, lumen-containing capillaries, and require supporting cells, such as fibroblasts or mesenchymal stromal cells^26,33^. We therefore analyzed RCC4 and RCC4-VHL spheroids cultured in either their standard medium (DMEM) or media derived from endothelial cell growth medium (ECGM2) or conditioned medium from human fibroblasts cultured in ECGM2 (NHDF-CM). After three days, RCC4 spheroids exhibited pronounced invasive behavior, characterized by numerous protrusions, reflected by high spheroid areas and perimeters, along with a low circularity index (**Figure 2B-C**). The addition of EGF to ECGM2 depleted of all growth factors (ECGM2-Δ5), ECGM2 depleted of VEGF (ECGM2-ΔVEGF), or NHDF-CM enhanced invasion compared to DMEM. Optical sections of spheroid centers revealed cell depletion within the core, surrounded by invading cells with elongated protrusions, notably in NHDF-CM (**Figure 2D**). In contrast, RCC4-VHL spheroids remained non-invasive, retaining a predominantly circular morphology, even in NHDF-CM, which induced only sparse, short protrusions (**Figure 2B-C**). Optical sections of RCC4-VHL spheroids displayed an organized structure surrounding a central cavity, forming a cyst-like architecture (**Figure 2D**). The cystic feature, evident in DMEM became more pronounced in NHDF-CM, where RCC4-VHL spheroids displayed thinner cell layer, a denser and more defined cortical actin ring, and a significantly enlarged central cavity (**Figure 2E**). These findings demonstrate that RCC4 spheroids exhibited EMT features with high invasive capacities, particularly in NHDF-CM. We successfully established culture conditions that preserved tumor-like characteristics of RCC4 cells, linked to the *VHL* status and that were compatible with the 3D model of vascularized microtumors.

### Fibroblast- and tumor cell-conditioned media differentially impact endothelial capillary morphogenesis and stability

To assess the compatibility of tumor and endothelial cells in 3D co-cultures, we first examined the impact of RCC4- or RCC4-VHL-conditioned medium on capillary network formation and stability, using a 3D collagen I hydrogel assay^28^. HUVECs were cultured in different conditioned media, using NHDF-CM as reference. Neither RCC4-nor RCC4-VHL-conditioned medium alone induced capillary formation. However, combining NHDF- and tumor cell conditioned media stimulated capillary network formation, albeit to a significantly lesser extent than NHDF-CM alone, as indicated by reduced capillary length, branch point number, and connectivity index, in 3D quantification analyses (**Figure 3A-B**). Furthermore, in this combined treatment with NHDF-CM, the RCC4-conditioned medium provided a less complex capillary network than RCC4-VHL-conditioned medium, as reflected by a significantly lower branch point number and connectivity index. These findings demonstrate that, although containing VEGF and promoting 2D migration of HUVECs (**Supplementary Figure S3A**) RCC4-condition medium was insufficient to support the formation of a complex capillary network, requiring the addition of NHDF-CM. Moreover, the combination of fibroblast and RCC4-conditioned media induced a capillary morphogenesis pattern distinct from that observed with RCC4-VHL-conditioned medium.

**Figure 3.**
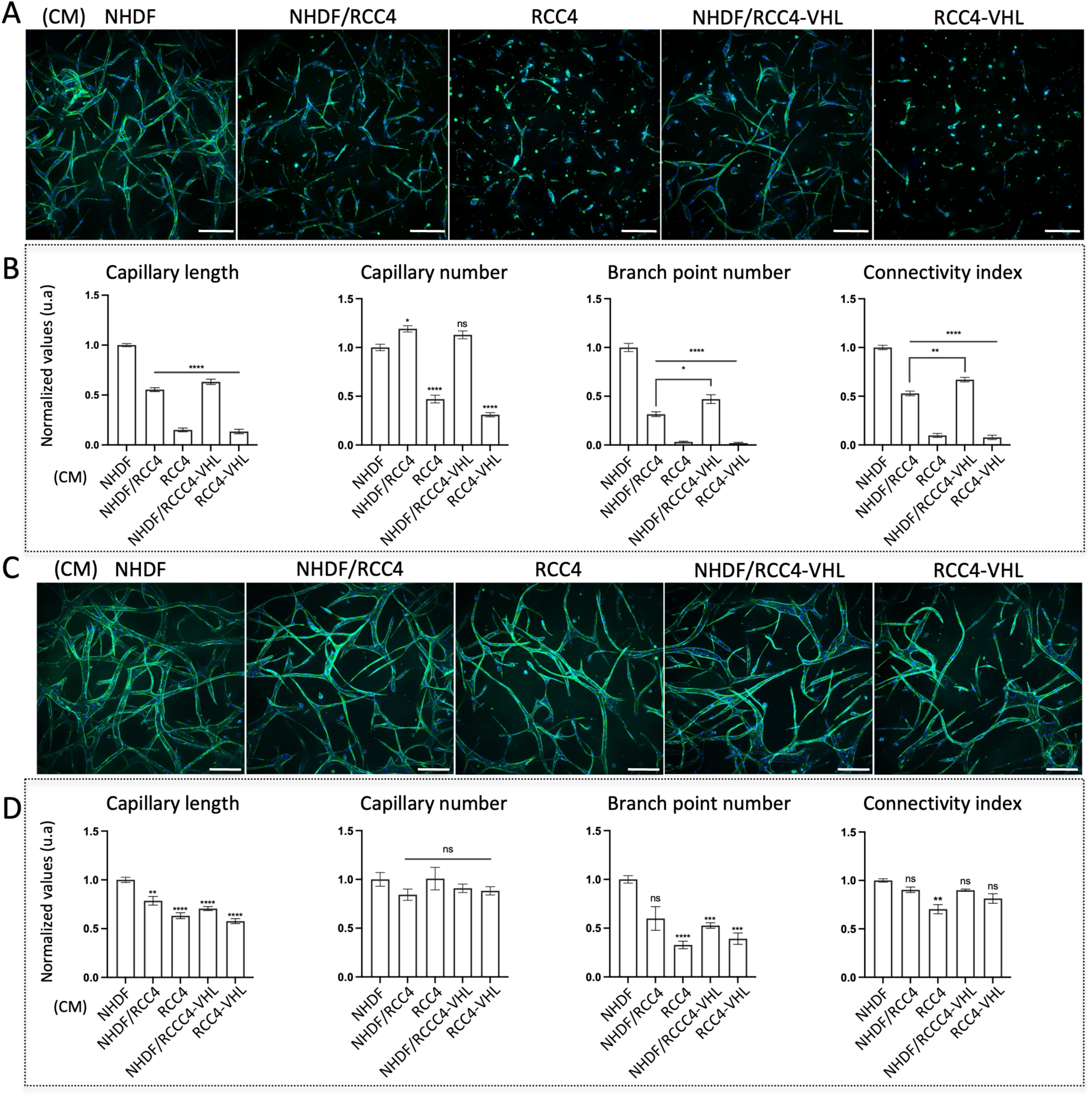
Conditioned medium from tumor cells alter capillary formation and stability in 3D models. A: Capillaries of HUVEC seeded in collagen I hydrogel and cultured for 5 days in conditioned medium (CM) from fibroblast (NHDF), RCC4, RCC4-VHL cells or 1:1 mixture of NHDF-CM/RCC4-CM or of NHDF-CM/RCC4-VHL-CM, stained for F-actin by phalloidin (green) and for nuclei by DAPI (blue). Images are displayed as maximum intensity projections of the capillary network (260 µm z-stacks). Scale bar: 200 µm. B: 3D quantification of total capillary length, capillary number, branch point number and connectivity index. Results were normalized to control condition (NHDF-CM). C: Capillaries formed and grown for 5 days in NHDF-CM were further treated for 4 days by the various cell-derived CM described in A. Images were performed as in A. Scale bar: 200 µm. D: 3D quantifications were performed as in B. Results were normalized to control condition in NHDF-CM. Graphs represent the mean of 3 independent experiments +/- SEM. Statistical analyses were performed using Brown-Forsythe and Welch ANOVA tests and represented with regard to the values of NHDF-CM (B and D). ns p>0.05, * p<0.05; ** p<0.01; *** p<0.001; **** p<0.0001.

A 3D angiogenesis assay^26^ provided additional valuable insights into capillary morphogenesis. Whereas NHDF-CM stimulated the formation of long, branched capillaries, RCC4-conditioned medium induced only 3D endothelial cell migration, without promoting capillary sprouting (**Supplementary Figure S3B-C**). Combining NHDF- and RCC4-conditioned media induced many capillary sprouts, although displaying fewer branch points, lower connectivity index than NHDF-CM (**Supplementary Figure S3B-C**) and discontinuous CD31 staining, forming empty sleeves of basement membrane (**Supplementary Figure S3D-E**). These findings highlighted that RCC4-conditioned medium negatively tuned capillary formation induced by CM-NHDF, and altered its morphogenesis.

To assess the stability of pre-formed capillary networks, mature capillaries generated in NHDF-CM were exposed to tumor-conditioned media (**Figure 3C**). When applied alone, RCC4- and RCC4-VHL-conditioned media disrupted the pre-existing networks. The destabilization effect was more pronounced when RCC4- or RCC4-VHL-conditioned media were applied alone compared to their combination with NHDF-CM. Moreover, these media differentially affected capillary connectivity and branching (**Figure 3D**). These observations suggest that RCC4 and RCC4-VHL cells negatively impact both the formation and stability of capillary networks, interfering with fibroblast-induced angiogenesis. Furthermore, these data suggest that RCC4 and RCC4-VHL differentially induced capillary morphogenesis, generating specific endothelial patterns.

### 3D vascularized microtumors mimic the ponds of the ccRCC vasculature

To investigate the influence of ccRCC tumor cells on capillary morphogenesis, we developed a 3D model of vascularized microtumors. Based on findings described above, maintaining NHDF-CM in the 3D model of vascularized microtumors was essential to support capillary formation. RCC4 or RCC4-VHL spheroids were co-embedded with HUVECs in a collagen I hydrogel, cultured for 4 to 7 days (**Figure 4A**) and analyzed by confocal microscopy using z-stack projections (**Figure 4B**) and optical sections (**Figure 4C**). Both RCC4 and RCC4-VHL spheroids exhibited invasive behavior, with endothelial cells significantly enhancing tumor cell migration. The RCC4-VHL spheroids displayed a markedly different morphology from their cyst-like structure observed in **Figure 2D**, appearing as an invasive mass with numerous protrusions extending from the core. The RCC4 spheroids generated long multicellular protrusions and extensive individual migration of cells dispersing far from the spheroid core. Although endothelial capillaries formed in close contact with both spheroid types, their networks exhibited distinct structural differences. Around RCC4-VHL spheroids, capillaries were sparsely distributed and poorly organized, whereas RCC4 spheroids accumulated capillaries at their core, forming a denser pattern (**Figure 4B**). 3D quantification revealed that RCC4 spheroids induced longer, more branched capillaries, with enhanced connectivity compared to RCC4-VHL spheroids (**Figure 4D**). Optical sections taken at different spheroid levels demonstrated that capillaries were in close contact with the outer surface of RCC4-VHL spheroids but did not penetrate the cell layers (**Figure 4C**), highlighting the cohesive and tightly organized nature of these spheroids. In contrast, small endothelial-lined cavities observed at the bottom and top levels of RCC4 spheroids converged into an irregular cavity settled in the central core (**Figure 4C** and **Supplementary Movie S7**). Remarkably, this inner cavity exhibited dimensions and shape resembling the pond identified in patient samples and PDX. However, this model still presents limitations, notably its inability to establish complete endothelial sheets. The pond structure within RCC4 spheroids was delineated by a polarized basement membrane closely associated with the basal side of the endothelial monolayer (**Figure 4E**). Higher magnification revealed a dense meshwork of collagen IV at the interface between endothelial and tumor cells. Unlike endothelial cells, tumor cells did not spread along the basement membrane but were packed around small collagen IV aggregates. The pond cavity contained isolated dying endothelial cells, displaying nuclear fragmentation, while the endothelial monolayer and surrounding tumor cells exhibited intact nuclei. These observations suggest that pond formation is a dynamic process involving homotypic interactions among endothelial cells and heterotypic interactions between endothelial and tumor cells, mediated by the basement membrane interface and accompanied by cell migration and selective survival. These findings demonstrate that the 3D co-culture model faithfully recapitulates the endothelial pond features of ccRCC, driven by the capacity of RCC4 spheroids to orchestrate endothelial cell organization and by tight tumor-endothelial cell interactions within a defined time-frame.

**Figure 4.**
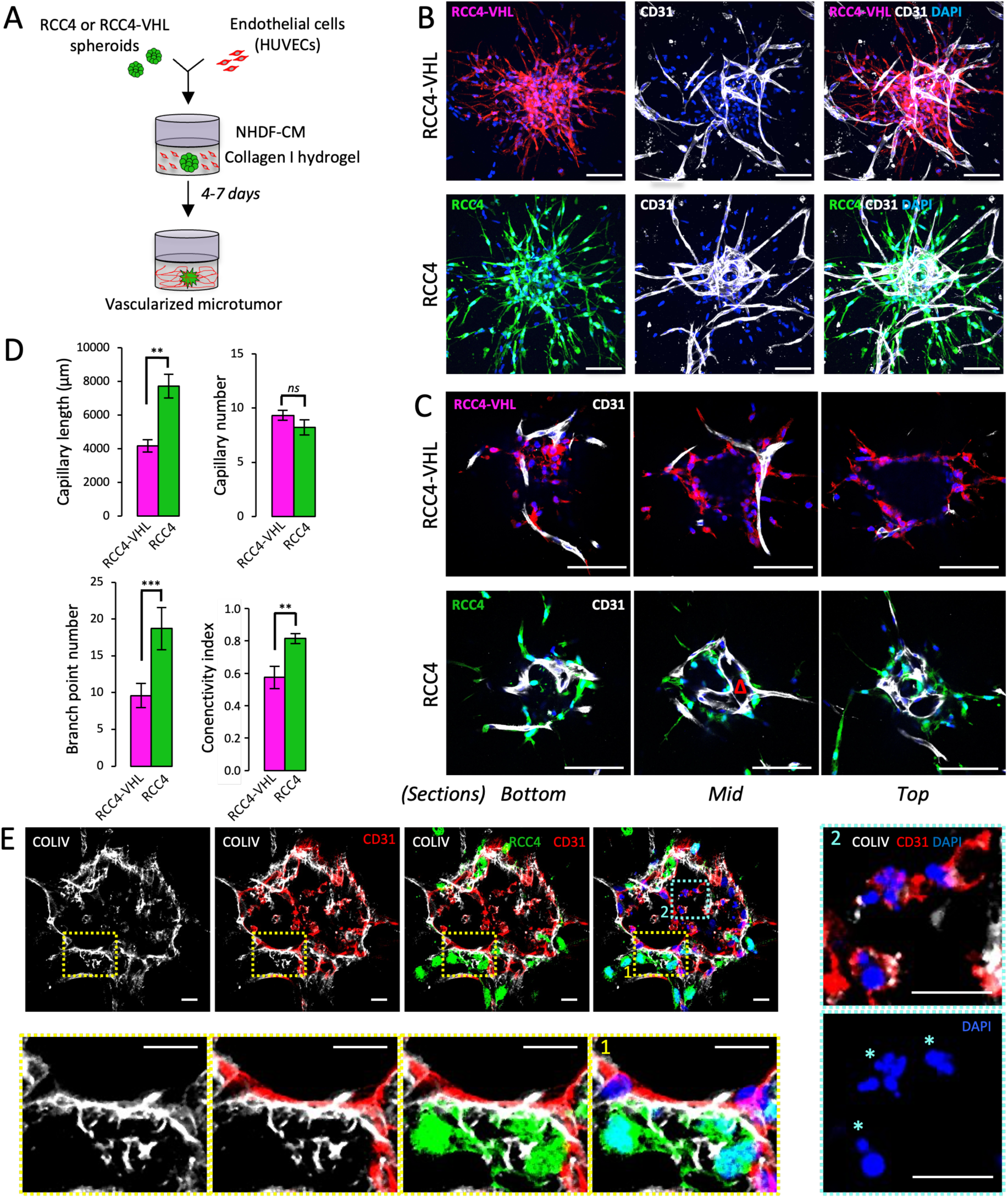
RCC4 spheroids increase capillary network and trigger pond formation in a 3D vascularized microtumor model. **A**: Schematic representation of 3D co-culture model. Spheroids of fluorescent RCC4 or RCC4-VHL were co-embedded in collagen I hydrogel with HUVEC suspension and cultured for 4 to 7 days in fibroblast-conditioned medium (NHDF-CM). **B-E**: Spheroids of fluorescent RCC4 (green) or RCC4-VHL (red) and capillaries after 4 days of culture. Images of vascularized microtumors immunostained for CD31 (white) and stained by DAPI for nuclei (blue) are displayed as z-stack projections (**B**) and optical sections at 3 spheroid z-levels (**C**). The endothelial pond is identified by a red triangle in the mid-plane section of RCC4 spheroid. Scale bar: 100 μm. 3D quantification of total capillary length, capillary number, branch point number and connectivity index of the capillary network formed in RCC4 (green) or RCC4-VHL (red) spheroid (**D**). Graphs represent the mean of 3 independent experiments +/- SEM. Statistical analyses were performed using Mann-Whitney test. ns p>0.05, ** p<0.01, *** p<0.001. Mid-plane section of RCC4 (green) spheroid reveals an endothelial pond immunostained for collagen IV (white), CD31 (red), and stained by DAPI for nuclei (blue) (**E**). Magnifications of yellow-(area 1) and blue-dotted (area 2) frames display the tumor-endothelial cell interactions at the pond periphery and fragmented nuclei (blue asterisk) of endothelial cells in the pond cavity, respectively. Scale bar: 20 μm.

### Dynamic evolution of pond morphogenesis in the 3D vascularized microtumor model

To further explore the dynamics of pond formation, time-lapse confocal imaging was conducted every 10–12 hours over 140 hours, using fluorescent endothelial cells and RCC4 or RCC4-VHL cells in the 3D co-culture model (**Figure 5**). Consistent with its weak invasive capacity, the RCC4-VHL spheroid exhibited slow protrusion elongation (**Figure 5A**). Endothelial cells, initially uniformly distributed within the hydrogel, gradually extended and fused, forming capillary segments up to 44 hours. From 68 hours, capillaries converged toward the spheroid, establishing direct interactions with its surface. In contrast, the rapid and extensive invasiveness of RCC4 cells resulted in spheroid scattering, rendering its initial volume undetectable by 20 hours. Between 20 and 44 hours, abundant multicellular protrusions formed, progressively replaced by the individual cell migration starting around 92 hours (**Figure 5B**). Meanwhile, the endothelial cells displayed a distinctly different dynamic behavior in the presence of RCC4 spheroids compared to RCC4-VHL spheroids. As early as 10 hours after incorporation into the hydrogel, elongated endothelial cells began to fuse and form growing capillary segments, toward the RCC4 spheroid by 20 hours. From 32 hours, converging capillary segments accumulated at the spheroid core, and fused by 44 hours. The endothelial network rapidly expanded, closely interacting with the invading tumor cells and connecting with numerous converging capillaries. Higher magnification of optical sections showed that endothelial-lined cavities within the scattering RCC4 spheroid underwent continuous remodeling, resulting into a unique pond, which progressively expanded until 116 hours (**Figure 5C**). Dying endothelial cells from regressing capillary segments were observed within the cavities, as seen in **Figure 4E**. Although the mature pond maintained a stable architecture between 116 and 140 hours, angiogenic process persisted. Notably, at 128 hours, a sprouting capillary, led by a migrating tip cell within the collagen I microenvironment, anastomosed with the adjacent pond wall by 140 hours (**Supplementary Figure S4**). Importantly, co-culturing endothelial cells with individual RCC4 cells dispersed in the hydrogel did not trigger pond formation (**Supplementary Figure S5**). Moreover, increasing the number of RCC4 cells in this co-culture model dramatically altered the capillary network, consistent with the destabilizing paracrine effect of RCC4 cells seen in **Figure 3**. These results highlight RCC4’s enhanced capacity to promote the formation of a complex and interconnected endothelial network through homotypic endothelial interactions and heterotypic tumor-endothelial interactions within a defined temporal and spatial framework. The vascularized microtumor model enables a sequential process, whereby tumor cells first induced endothelial network formation through paracrine signaling before directly interacting with converging capillaries, thereby promoting pond morphogenesis.

**Figure 5.**
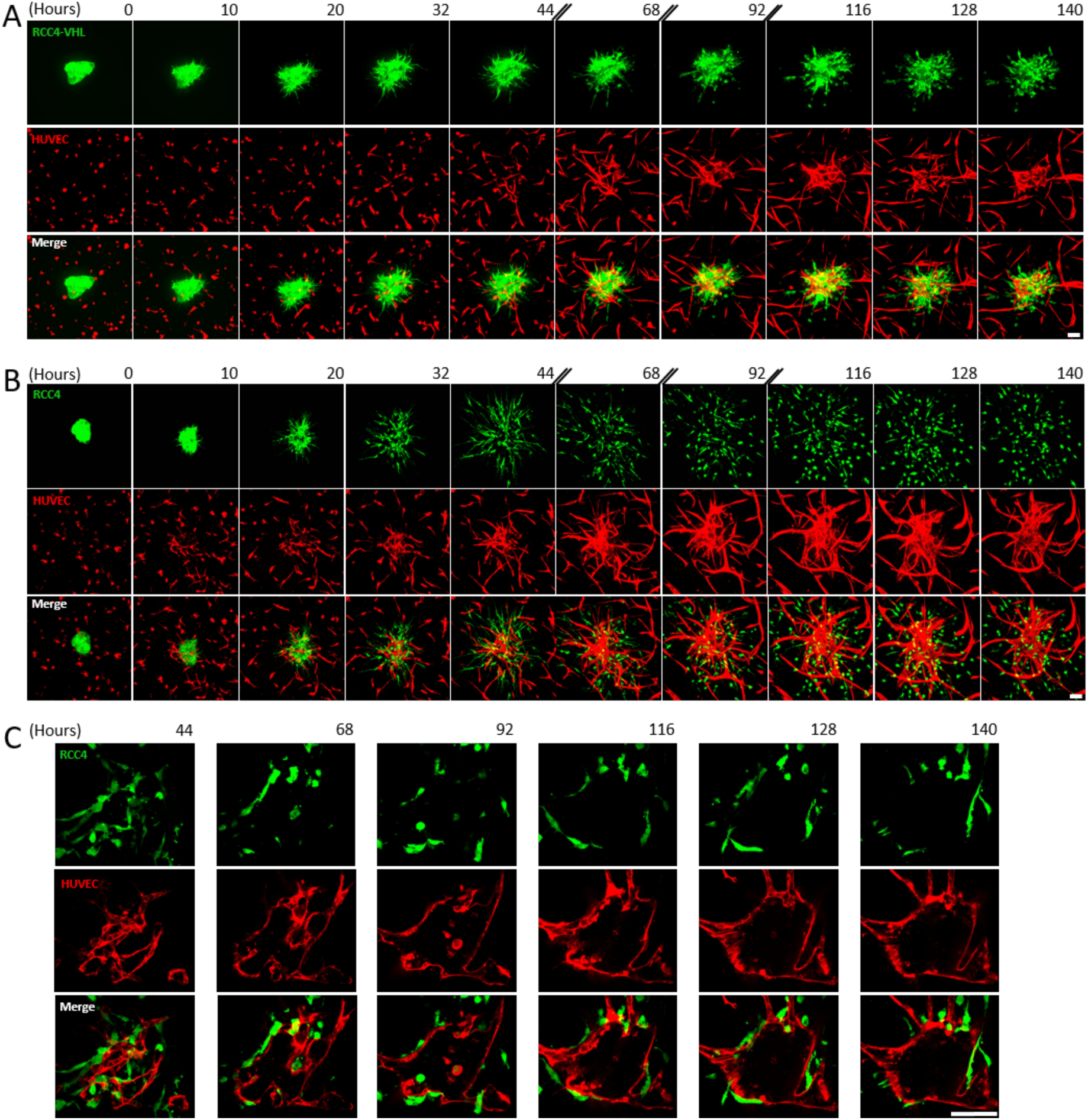
Pond establishment requires converging endothelial sprouts towards the tumor spheroid and close interactions between the RCC4 cells and endothelium in the vascularized microtumors. A and B: Time-lapse images of vascularized spheroids of RCC4-VHL (A) or RCC4 (B). Fluorescent spheroids (green) were co-embedded with fluorescent HUVEC (red) in collagen I hydrogel and cultured for 6 days in fibroblast-conditioned medium. Images of the spheroid were acquired every 10 to 12 hours and are displayed as maximal z-stack projections. Scale bar: 50 µm. C: Formation and remodeling processes of the endothelial pond in the invading RCC4 spheroid. Images were acquired between 44 and 140 hours of culture and are displayed for optical z-mid-plane sections. Scale bar: 100 µm.

### Ponds, tumor capillaries and tumor cells exhibit distinct sensitivities to therapeutic treatments

While previous studies have explored the effects of temsirolimus^34^, crizotinib^2^, and sunitinib^35,36^ on apoptosis, proliferation of ccRCC cells or spheroids and capillaries in 2D or in *in vivo* models, their impact on tumor invasiveness and capillary morphogenesis within a 3D microenvironment remains poorly explored. To address this gap, we evaluated the effects of the three inhibitors on capillary network formation, tumor cell invasion, and pond morphogenesis within the vascularized microtumor model. After a 2-day treatment of endothelial cells (**Figure 6A-B**), RCC4 spheroids (**Figure 6C-D**), or both (**Figure 6E-G**), embedded into collagen I hydrogel, we analyzed the capillary network using 3D quantification (**Figure 6B** and **F**), and the tumor invasion by measuring spheroid area and perimeter (**Figure 6D** and **G**). Temsirolimus, at all tested concentrations, significantly reduced capillary length, branching, and connectivity (**Figure 6A-B**), as well as spheroid size and perimeter (**Figure 6C-D**), highlighting its strong anti-angiogenic and anti-invasive effects. Crizotinib exhibited a dose-dependent reduction in spheroid area and perimeter (**Figure 6C-D**) and to a lesser extent on capillary length and connectivity index (**Figure 6A-B**), indicating moderate anti-invasive and anti-angiogenic properties. Sunitinib significantly decreased capillary length and connectivity across all concentrations (**Figure 6A-B**), while reducing spheroid area and perimeter at the two highest concentrations (**Figure 6C-D**). These dose-response profiles suggest that sunitinib primarily modulated capillary morphogenesis, while temsirolimus affected both capillary formation and tumor spheroid invasion, and crizotinib predominantly targeted tumor invasion.

**Figure 6.**
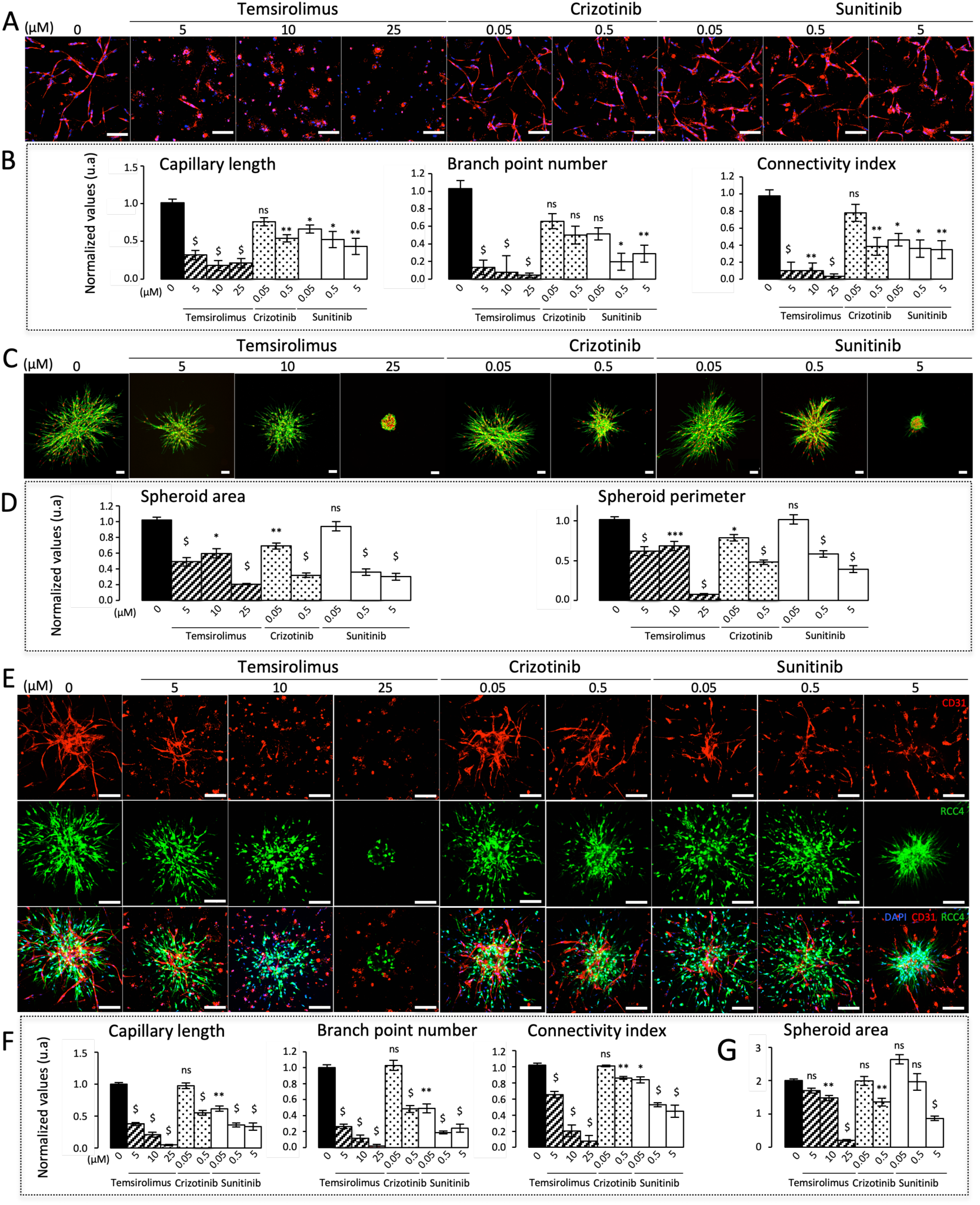
Therapeutic treatments against ccRCC differentially inhibit tumor invasiveness and capillary formation in vascularized microtumors. HUVEC (A and B), RCC4 spheroids (C and D) or co-culture of HUVEC and RCC4 spheroids (E-G) were embedded in collagen I hydrogel and treated by temsirolimus (5, 10 or 25 µM), crizotinib (0.05 or 0.5 µM), sunitinib (0.05, 0.5 or 5 µM) or vehicle (0) in NHDF-conditioned medium for 2 days. A: Capillaries immunostained for CD31 (red) and stained bt DAPI for nuclei (blue). Scale bar: 100 µm. B: 3D quantification of total capillary length, branch point number and connectivity index of capillary network. Results were normalized to control condition (vehicle). C: RCC4 spheroids stained for F-actin by phalloidin (green) and for nuclei by DAPI (red). Scale bar: 100 µm. D: Quantification of area and perimeter of spheroids. Results were normalized to control condition (vehicle). E: Vascularized fluorescent RCC4 spheroids (green, mid- and bottom panels) immunostained for CD31 (red, top and bottom panels) and stained by DAPI for nuclei (blue, bottom panels). Scale bar: 100 µm. F: 3D quantification of total capillary length, branch point number and connectivity index of the capillary network in the vascularized microtumors. Results were normalized to control condition (vehicle). G: Quantification of area and perimeter of spheroids. Results were normalized to control condition (vehicle). Graphs represent the mean of 3 independent experiments +/- SEM. Statistical analyses were performed over control condition using Kruskal-Wallis test (B, D, F and G). ns p>0.05, * p<0.05, ** p<0.01, *** p<0.001, $ p<0.00001.

The presence of endothelial cells in the 3D co-culture model slightly reduced the anti-angiogenic efficacy of all three agents, shifting the inhibitory effects toward higher concentrations, particularly regarding the connectivity index (**Figure 6E-F**). he inhibition of spheroid invasion was more markedly attenuated, requiring the highest concentrations of all three drugs to achieve a significant effect (**Figure 6E** and **G**). These findings indicate that endothelial cells confer a degree of resistance to RCC4 cells against temsirolimus, crizotinib, and sunitinib in the 3D microenvironment.

Given these results, we assumed that the two lowest concentrations of sunitinib would affect the capillary network without significantly altering tumor spheroid invasion in the co-culture model. We then evaluated the sensitivity of pre-existing ponds and tumor capillaries to sunitinib at 0.05 or 0.5 µM, doses aligned with pharmacokinetic and pharmacodynamic analyses determining an efficient range between 0.125– 0.25 μM^37^. Consistent with the mature pond observed from 116 hours in **Figure 5C**, sunitinib treatment was initiated on day 5 of culture when vascularized microtumors contain a mature pond connected to converging capillaries (**Supplementary Figure S6**). Adjacent capillaries were disconnected from the pond and located in the adjacent plane of the same culture well. Vehicle or sunitinib (0.05 or 0.5 µM) was applied for 2 days (**Figure 7A**). 3D imaging revealed that pond architecture remained unaffected by sunitinb treatment at both concentrations (**Supplementary Movies S8-S10**). Vascular area quantification demonstrated a significant, dose-dependent regression of adjacent capillaries, while the ponds and their connected capillaries remained largely insensitive to the drug (**Figure 7B**).

**Figure 7.**
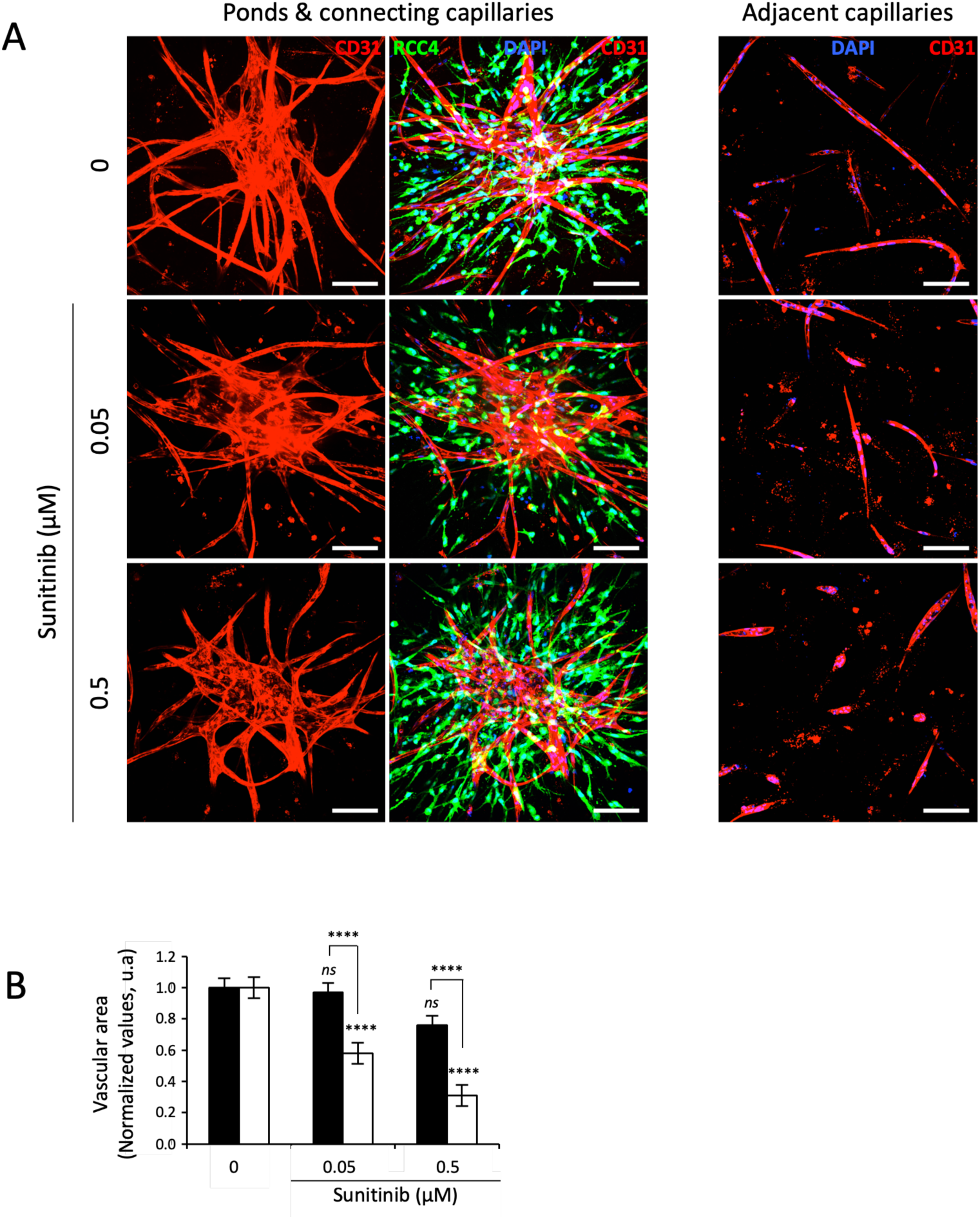
Endothelial ponds exhibit a low sensitivity to sunitinib in vascularized microtumors. **A**: vascularization of spheroids was promoted for 5 days in the presence of NHDF-CM after which treatment with sunitinib (0.05 or 0.5 µM) was applied for 2 days. Vascularized fluorescent RCC4 spheroids (green, mid-panels) immunostained for CD31 (red) and stained by DAPI for nuclei (blue, mid- and right panels). Maximal z-stack projection of spheroid images (left and mid-panels) display the pond and connecting capillaries. Images of the spheroid-adjacent field (right panel) display capillary networks. Scale bar: 100 µm. **B**: Quantification of vascular areas of pond and connecting capillaries (black columns) and of adjacent tumor capillaries (white columns) were performed on z-stack projections. Results were normalized to corresponding control condition (vehicle). Graphs represent the mean of 3 independent experiments +/- SEM. Statistical analyses were performed using ANOVA test. Statistical mentions above columns proceed from comparison of treated pond and converging capillaries or of treated adjacent tumor capillaries to their corresponding control condition. ns p>0.05, **** p<0.0001.

In conclusion, our 3D *in vitro* model, which accurately mimics the pond architecture observed in ccRCC patients, provided valuable insights into the differential sensitivities of tumor-associated vascular structures to therapeutic interventions. These findings demonstrate that the ponds exhibited lower sensitivity to sunitinib compared to the adjacent capillaries, suggesting that the presence and distribution of aberrant vascular structures in ccRCC patients could significantly influence treatment efficacy.

## DISCUSSION

ccRCC is a highly vascularized carcinoma, displaying a prominent CD31^+^ network in the tumor mass. Sunitinib, the first TKI that improved survival for patients with metastatic ccRCC, became the main-line treatment for nearly two decades and the reference treatment in subsequent clinical trials^17^. However, this anti-angiogenic treatment has reached its limits as many patients displayed either inherited resistance or developed resistance. We report here that the ccRCC vascular network contains aberrant endothelial structures which exhibit striking 3D architecture and dimensions, strongly differing from tumor capillaries: ponds are dilated and irregular structures containing a broad lumen; sheets are collapsed extended structures. These structures exhibit various distribution and abundancy in primary tumors from metastatic ccRCC. We set up a 3D *in vitro* model of vascularized microtumors able to recapitulate the endothelial pond exhibiting similar features than those of patients. Our data revealed that paracrine regulation and/or direct crosstalks between endothelial and ccRCC cells lead to pond formation and tuned sensitivity to therapeutic treatments. Our results suggest that ponds contribute to the sunitinib resistance and patient outcome.

The complexity of the ccRCC vascular network has been described for decades, based on the 2D assessment of tissue sections. In addition to the various vessel morphologies reported in many tumors, including breast cancer^38^ and hemangioblastoma^39^, careful examination of several previous studies reveals 2D structures in ccRCC patient samples that resemble ponds^40^ or ponds and sheets^39^. Yao and coll.^41^ displayed immunohistochemically stained sections of ccRCC presenting a high density of CD31^+^ or CD34^+^ structures similar to the ponds and sheets characterized in the present study. Ruiz-Sauri and coll.^8^ identified two types of CD31^+^ patterns present in ccRCC and absent in papillary and chromophobe renal cell carcinomas: the *pseudoacinar* one formed by round or elongated vessels with large lumens which, occasionally, are narrowed, correspond to the pond 2D architecture; the *fascicular* one, presenting elongated vessels with either narrow or collapsed lumens, correspond to the sheet 2D architecture. Ponds and sheets interconnect, suggesting a complex dynamic remodeling of the ccRCC vascular network and raising the question of endothelial network remodeling throughout tumor evolution.

Contradictory results have been reported regarding the potential correlation between ccRCC grades or patient outcome and presence of distinct endothelial patterns^6,8,41^, therefore requiring further clinical investigations. However, due to the high complexity of the ccRCC vascular network, obtaining its complete picture within the TME using a 2D approach is inadequate. Recently, Kaneko and coll.^42^ reported a 3D analysis of the ccRCC microvasculature after performing tissue clearing and volumetric immunohistochemistry. The authors claimed that the spatial elucidation of the volumetric vasculature could be prognostic and may serve as biomarker for genomic alterations, as they detected potential association of the PI3K-mTOR pathway with 3D vascular features, and with vessel radius. Although invaluable, this innovative approach is clinically and technically challenging for deciphering the cellular processes leading to the 3D architecture of the aberrant endothelial structures in ccRCC.

Our study combines two complementary approaches, the 3D imaging providing unprecedented insights into the architecture of ponds and sheets and an innovative vascularized microtumor model that successfully recapitulates ponds, exhibiting features similar to those observed in patients. Our current model fails to recreate sheets, raising the question of whether ponds gradually remodel into sheets or whether they are formed through two distinct mechanisms. However, the 3D vascularized microtumor model unveils several key features of human ccRCC, such as tumor-endothelial cells interactions, cell-ECM interactions, and paracrine regulation by fibroblasts contributing to the pond formation.

Tumor cells are key players for developing this pertinent model. We show here that RCC4 cells exhibit on the one hand, EMT features, consistent with previous studies^43,44^ in both 2D and 3D cultures, and on the other hand, EGF-induced invasive capacities, in line with the well-known activating mutations of the EGF receptor in ccRCC^45^. Furthermore, 3D culture of RCC4 cells in collagen I hydrogel more accurately recapitulate the gene expression profile of primary RCC and exhibit an enrichment of genes involved in EMT, compared to 2D culture^46^. Our study shows that RCC4 cells exhibiting ccRCC tumor-like features differentially affect capillary morphogenesis compared to RCC4-VHL cells and that ccRCC tumor cells undergoing EMT instruct endothelial cells, leading to specific capillary morphogenesis and pond formation. Similarly, in the PDX model, the human tumor cells instruct the mouse endothelial cells, resulting in the preservation/formation of ponds and sheets, consistent with the rapid replacement of human stromal components by the murine TME after transplantation^29,47^. Tumor cell-mediated instruction of endothelial cells occurs through both paracrine signaling and heterotypic cell-cell interactions, with fibroblasts tuning this process through paracrine effects. RCC4-conditioned medium not only fails to induce capillary formation, but also disrupts pre-existing capillaries and induces endothelial cell migration. The instructional process, carried out by ccRCC tumor cells may involve extracellular vesicules, as demonstrated by Takeda and coll, using the 786-O cell line^48^ or the secretion of VEGF, interleukin-8, and stromal cell-derived factor 1 (SDF-1) mediated by the low expression of monocyte chemotactic protein 1–induced protein 1 as reported by Marona and coll. using Caki-1/-2 cell lines^40^. Zagzag and coll.^39^ have shown in patients and RCC4 cell line that VHL loss-of-function leads to increased expression of both CXCR4 and its ligand SDF-1, establishing paracrine and autocrine signaling pathways that may contribute to ccRCC exuberant vascularization. Beyong the impact of *VHL*-mutated tumor cells on endothelial cells, the enhanced invasion of RCC4 cells, an EMT hallmark, triggered by the endothelial cells unveils the complex, reciprocal crosstalk between the two constituent cells within the TME. These findings align with previous work demonstrating that *VHL*-deficient cells reprogram endothelial cells which in turn facilitate tumor cell invasion through oncostatin M secretion^49^. In addition, our study emphasizes the cooperative involvement of tumor-and fibroblast-conditioned media in capillary morphogenesis, mimicking the paracrine regulation of peri-tumoral capillaries by the tumor mass. As converging capillaries contact the invading tumor, the endothelial network anastomoses, as indicated by the high branch point number and connectivity index, forming small cavities, that eventually fuse to form the pond. Meanwhile, capillary destabilization induced by the RCC4-conditioned medium contributes to pond lumen formation. This 3D approach highlights the temporal and spatial dynamics of paracrine and direct reciprocal interactions, driving the specific morphogenesis required for pond formation. The complexity and multiplicity of these cellular interactions participate to the intra- and inter-heterogeneity in the abundance and distribution of aberrant vascular structures in ccRCC tumors.

We hypothezise that the aberrant vascular structures contribute to the partial and variable efficacy of targeted therapies for metastatic ccRCC. The response to treatments integrates the complex interplay between cells or between cells and ECM components within a 3D microenvironment. Several models have emerged for drug assessment, including PDX, whose advantages and limitations have been mentioned above, as well as organoid models. The patient-derived renal cell carcinoma organoid model has shown promising results in establishing differential sensitivity patterns based on intertumoral heterogeneity^50^. However, this model still faces limitations regarding long-term co-culturing capabilities for multiple cell types, varying proportions of endothelial cells in organoids from different patients, and an inability to undergo endothelial network morphogenesis. The present vascularized microtumor model provides a suitable research platform for assessing TKI treatments, as it faithfully recapitulates ponds. We demonstrate, on the one hand, that endothelial cells decrease tumor cell sensitivity to temsirolimus, crizotinib and sunitinib and enhance their invasive capacities, and on the other hand, that ponds are less sensitive to sunitinib than adjacent tumor capillaries.

The main issue is now to further understand the molecular mechanisms underlying the low sensitivity to sunitinib as well as discovering new drugs able to target the resistant vascular structures. Works using single cell-RNA sequencing performed on ccRCC samples revealed two major endothelial clusters, one exhibiting higher expression of VEGF receptors 1 and 2 mRNAs compared with the other^51–54^. Authors claimed that the second endothelial cluster may escape angiogenesis inhibitors. We therefore suggest that the pond is composed of endothelial cells of the second cluster, providing a decreased sensitivity to sunitinib.

Consistent with our *in vitro* data demonstrating the low sensitivity of ponds to sunitinib compared to tumor capillaries, a careful analysis of sunitinib-resistant tumors in PDX model, unveils numerous ponds whereas sunitinib-sensitive ones exhibit sparse capillaries^55^. In addition to its direct effect on the endothelial network, sunitinib treatment impacts tumor cells. Metastatic ccRCC tumor samples before treatment and immediately after disease progression to a TKI. Microarray analysis of metastatic ccRCC primary tumors pre- and post-treated by TKI, including sunitinib, demonstrated an increased expression of EMT-related genes in TKI-resistant cells, acquiring migration and invasion capacity^56^. Taken together, these data suggest that sunitinib can increase the pond density versus the tumor capillary density through their differential sensitivity and through the increased EMT undergone by the TKI-resistant tumor cells. This closed loop mechanism involving tumor plasticity and endothelial cell instruction may lead to ponds formation and contribute to the aggressive behavior of TKI-resistant ccRCC.

Altogether we characterized the 3D architecture of the intratumoral aberrant vascular structures of ccRCC, either dilated for the ponds or collapsed for the sheets. We provided an innovative and relevant approach of 3D vascularized microtumor model favoring the paracrine or direct interactions between fibroblasts, endothelial cells and tumor cells in an ECM microenvironment. These approaches allowed us to recapitulate the ccRCC endothelial pond and to demonstrate its low sensitivity to sunitinib. This model provides a unvaluable tool for assessing the responses of ccRCC to targeted treatments. This study opens new avenues for disrupting endothelial ponds leading to the decrease of both perfusion of the local intratumoral mass and the metastatic spread of aggressive ccRCC.

## Author contributions

N.B-J., Y.A., C.C., L.M. and C.M. conceived the study, designed experiments, conducted experiments and interpreted results; C.A-R., M.E.B., N.J., C.L., S.D.O and C.H. conducted experiments; P.M. interpreted results; V.L. and G.B. conducted experiments and analyzed data; I.C., M.S., G.B. and S.G. interpreted results; N.B-J., Y.A. and C.M. wrote the manuscript.

## Acknowledgements

We gratefully acknowledge the Orion technological core (IMACHEM-IBiSA) of Center for Interdisciplinary Research in Biology for their support, and especially Estelle Anceaume for access to the microtome, Julien Dumont and Tristan Piolot for assisting us with slide scanner acquisition and Héloïse Monnet and Philippe Mailly for developing image analysis scripts presented in this article. We thank members of AP-HP, Hôpital Saint-Louis, Unité de Thérapie Cellulaire, CRB-Banque de Sang de Cordon, Paris, France, and especially Thomas Domet for providing human umbilical cords. Noémie Brassard-Jollive was supported by Sorbonne Université and by Fondation ARC Jeunes Chercheurs. Yoann Atlas was supported by La Ligue Contre le Cancer. The work was supported by the Gefluc and by the Fondation ARC (PJA 20161205085).

