## Supplemental figures 1-6 for "3D vascularized microtumors unveil aberrant ccRCC vasculature and differential sensitivity to targeted treatments"

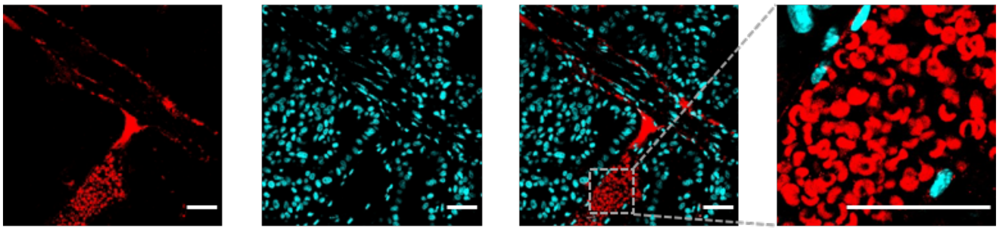

**Supplementary Figure S1. Lumen of endothelial pond containing red blood cell aggregates.**
500 µm-thick section of ccRCC-derived xenograft immunostained for endothelium by anti-CD31 antibody (red), stained for red blood cells by auto-fluorescence and for nuclei by DAPI (blue). Scale bar: 50 µm.

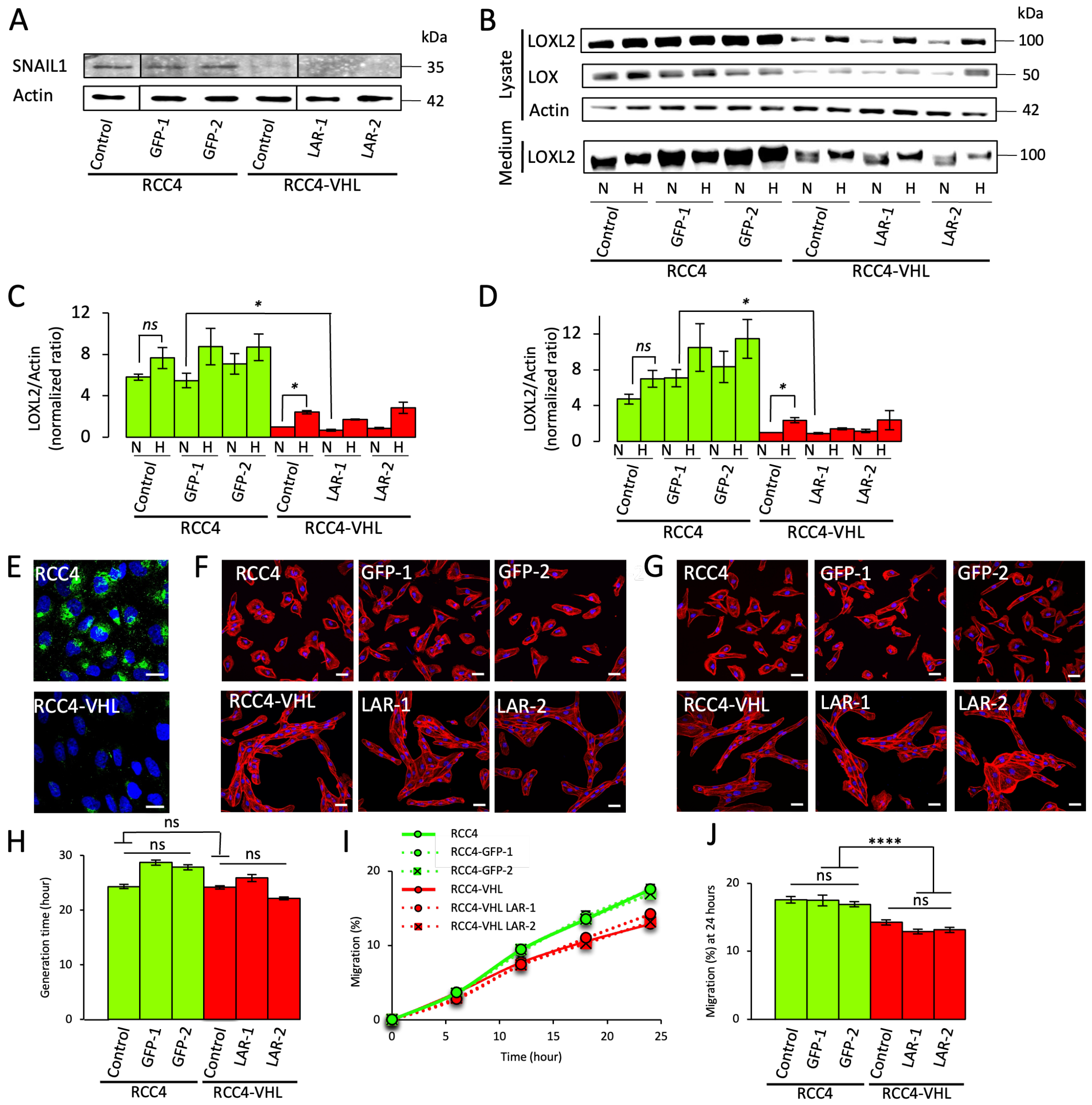


**Supplementary Figure S2. Fluorescent clones of RCC4 or RCC4-VHL exhibit similar features to their parental cell line.**

EMT properties were investigated for parental RCC4 cell line (RCC4) and corresponding GFP-expressing clones (GFP-1 and -2) or for parental VHL-expressing RCC4 cell line (RCC4-VHL) and corresponding LAR-expressing clones (LAR-1 and -2). A-D: Cells were cultured for 2 days in normoxia (A) and in normoxia (N) or hypoxia corresponding to 1% O2 (H) (B-D). Proteins were extracted from cell lysate and analyzed by immunodetection for SNAIL1 expression (A) or from cell lysate and secretion medium and analyzed by immunodetection for LOXL2 and LOX amount (B). Actin was used as a loading control. Ratio of LOXL2 over actin amounts was quantified in the cell lysate (C) and secretion medium (D) from 3 independent experiments performed in duplicate. E: RCC4 and RCC4-VHL cells were cultured in 2D for 3 days and immunostained for LOXL2 (green). Nuclei (blue) were stained by DAPI. Scale bar: 10 µm. F-G: RCC4 and RCC4-VHL cell lines or clones were cultured for 2 days on plastic (F) or collagen I coating (G) and stained for F-actin by phalloidin (red) and nuclei by DAPI (blue). Scale bar:

50 µm. H-J: Proliferation and collective migration were monitored by live-cell imaging. Proliferation was assessed by generation time determined after 96 hour-culture (H). Migration was quantified over 24 hour-culture (I) and endpoints were plotted at 24 hour (J). Graphs represent the mean of 3 independent experiments ± SEM. ns $p > 0.05$ (Kruskal-Wallis test) (H) and ns $p>0.05$, **** $p<0.0001$ (one-way ANOVA) (J).

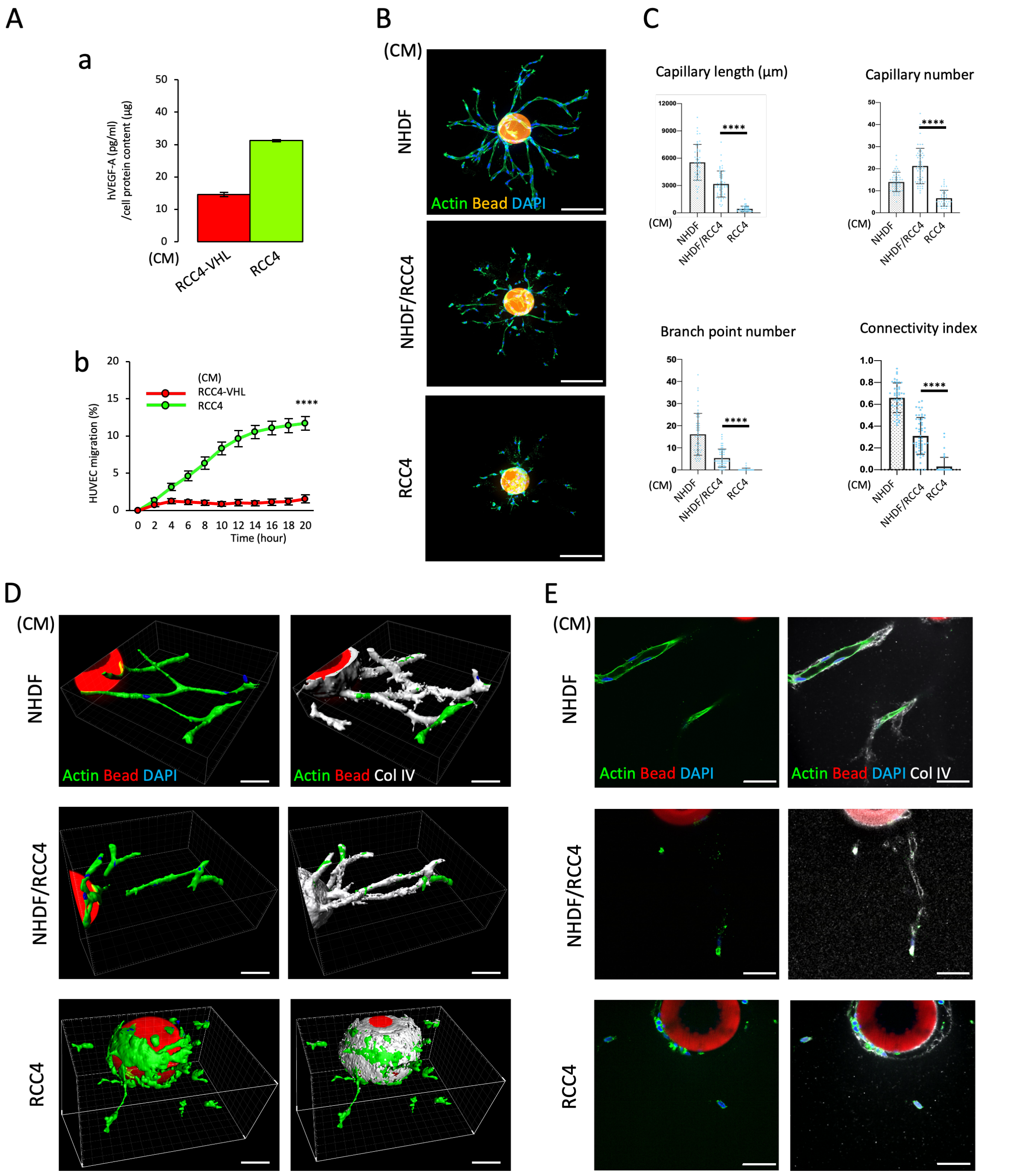


**Supplementary Figure S3. RCC4 conditioned medium alter capillary morphogenesis.**

A: Conditioned media (CM) of RCC4 or of RCC4-VHL cells were analyzed for their VEGF amount (a) and ability to induce endothelial cell 2D-migration (b). HUVEC migration was monitored over 20 hours of culture by live-cell imaging. Graphs represent the mean of 3 independent experiments ± SEM. **** p<0.0001 (One-Way ANOVA). B-E: Angiogenesis assay using fluorescent Cytodex beads covered by HUVEC monolayer embedded in fibrin hydrogel and cultured for 4 days in conditioned medium (CM) of fibroblast (NHDF) or RCC4 cells or a 1:1 mixture of CM (NHDF/RCC4). Images of capillary sprouts stained for F-actin by phalloidin (green) and nuclei by DAPI (blue) are displayed as z-stack projections. Scale bar: 200 µm (B). 3D quantifications of total capillary length, capillary number, branch point number and connectivity index. Graphs represent the mean of 3 independent experiments +/- SEM. **** $p<0.0001$ (Kruskal-Wallis test) (C). Images of capillary sprouts stained for CD31 (red), DAPI (blue) and collagen IV (white) are displayed as 3D reconstruction (D) or optical section (E). Scale bar: 50 µm.

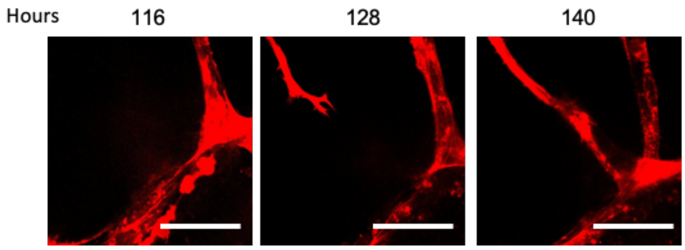


**Supplementary Figure S4. Angiogenic capillary sprouts towards the pond and connects to its endothelium.**

Fluorescent HUVEC (red) are displayed in magnified images of the vascularized RCC4 spheroid acquired between 116 and 140 hours of culture. Scale bar: 50 µm.

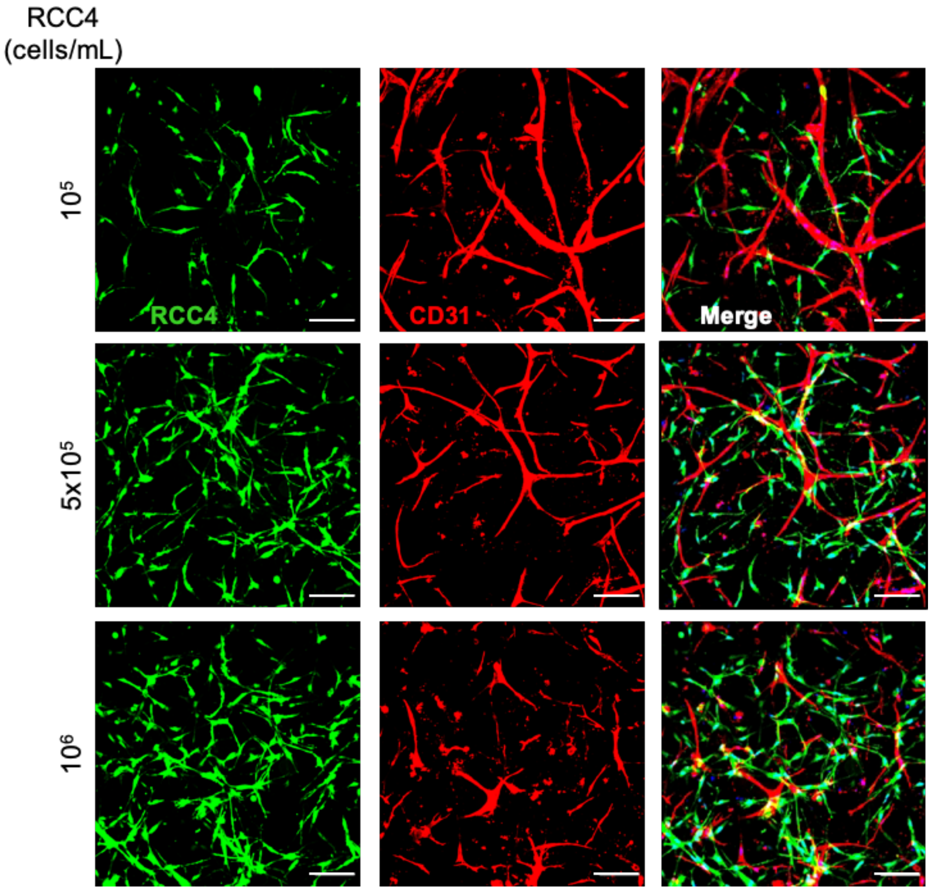


**Supplementary Figure S5. Tumor spheroid initial architecture is required for the pond formation.** Individualized RCC4 cells seeded at 106, 5x105 or 105 cells/mL and endothelial cells (1.5x106 cells/mL) were co-cultured in collagen I hydrogel for 5 days in presence of NHDF-CM. Images of fluorescent RCC4 (green), endothelial cells immunostained for CD31 (red) and nuclei stained by DAPI (blue) were displayed as z-stack projections. Scale bar: 100 µm.

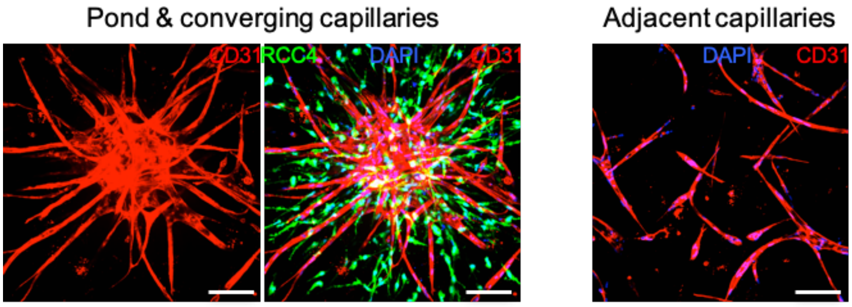


**Supplementary Figure S6. Vascularized RCC4 microtumors cultured for 5 days allowing formation of capillaries and pond.**

Vascularized fluorescent RCC4 spheroids (green, mid-panels) immunostained for CD31 (red) and stained for nuclei by DAPI (blue, mid-and right panels). Maximal z-stack projection of spheroid images (left and mid-panels) display the pond and converging capillaries. Images of the spheroid-adjacent field (right panel) display capillary networks. Scale bar: 100 µm.

**Supplementary Movie S1. Aberrant ccRCC vasculature in patient-derived xenograft of ccRCC analyzed by surface rendering and 3D reconstruction.**

Image tiles from 500 µm-thick section of xenograft sample were stitched. Movie displays 3D reconstructions of the aberrant vasculature stained for CD31 (red) surrounded by the cell mass stained by DAPI (blue) followed by a moving clipping plane performed on the volume rendering (Imaris software). The initial stage of the 3D imaging clearly revealed an isolated capillary located behind the main pond.

**Supplementary Movie S2. Red blood cells accumulated in pond of patient-derived xenograft of ccRCC analyzed by z-stack imaging.**

Movie displays z-stack images of 500 µm-thick section of xenograft stained for CD31 (red) and nuclei (blue). Red blood cells were autofluorescent (red).

**Supplementary Movie S3. Formation of the RCC4 spheroid monitored by transmitted-light time-lapse acquisitions.**

Movie displays images of 500 RCC4 cells seeded in low-attachment U-bottom plates and monitored for 9 hours.

**Supplementary Movie S4. Formation of the RCC4 spheroid monitored by fluorescent time-lapse acquisitions.**

Movie displays images of 500 RCC4 cells seeded in low-attachment U-bottom plates and monitored for 9 hours.

**Supplementary Movie S5. Formation of the RCC4-VHL spheroid monitored by transmitted-light time-lapse acquisitions.**

Movie displays images of 500 RCC4 cells seeded in low-attachment U-bottom plates and monitored for 9 hours.

**Supplementary Movie S6. Formation of the RCC4-VHL spheroid monitored by fluorescent time-lapse acquisitions.**

Movie displays images of 500 RCC4 cells seeded in low-attachment U-bottom plates and monitored for 9 hours.

**Supplementary Movie S7. Vascularized RCC4 microtumor analyzed by surface rendering and 3D reconstruction.**

Vascularized RCC4 spheroid was cultured for 5 days in collagen I hydrogel. Movie displays 3D reconstructions of CD31 (red) staining and fluorescent RCC4 (green) signal followed by a moving clipping plane performed on the volume rendering of the CD31 staining (Imaris software).

**Supplementary Movie S8. Vehicle-treated pond analyzed by surface rendering and 3D reconstruction.**

Vascularization of the RCC4 spheroid was promoted for 5 days in collagen I hydrogel in the presence of NHDF-CM after which treatment with vehicle was applied for 2 days. Movie displays 3D reconstruction performed on the volume rendering of the CD31 staining (red) (Imaris software).

**Supplementary Movie S9. Sunitinib (0.05µM)-treated pond analyzed by surface rendering and 3D reconstruction.**

Vascularization of the RCC4 spheroid was promoted for 5 days in collagen I hydrogel in the presence of NHDF-CM after which treatment with Sunitinib (0.05µM) was applied for 2 days. Movie displays 3D reconstruction performed on the volume rendering of the CD31 staining (red) (Imaris software).

**Supplementary Movie S10. Sunitinib (0.5µM)-treated pond analyzed by surface rendering and 3D reconstruction.**

Vascularization of the RCC4 spheroid was promoted for 5 days in collagen I hydrogel in the presence of NHDF-CM after which treatment with Sunitinib (0.5µM) was applied for 2 days. Movie displays 3D reconstruction performed on the volume rendering of the CD31 staining (red) (Imaris software).
